# DNA-m6A calling and integrated long-read epigenetic and genetic analysis with fibertools

**DOI:** 10.1101/2023.04.20.537673

**Authors:** Anupama Jha, Stephanie C. Bohaczuk, Yizi Mao, Jane Ranchalis, Benjamin J. Mallory, Alan T. Min, Morgan O. Hamm, Elliott Swanson, Danilo Dubocanin, Connor Finkbeiner, Tony Li, Dale Whittington, William Stafford Noble, Andrew B. Stergachis, Mitchell R. Vollger

**Author notes:** These authors contributed equally to this work (Anupama Jha, Stephanie C. Bohaczuk).

## Abstract

Long-read DNA sequencing has recently emerged as a powerful tool for studying both genetic and epigenetic architectures at single-molecule and single-nucleotide resolution. Long-read epigenetic studies encompass both the direct identification of native cytosine methylation as well as the identification of exogenously placed DNA *N^6^*-methyladenine (DNA-m6A). However, detecting DNA-m6A modifications using single-molecule sequencing, as well as co-processing single-molecule genetic and epigenetic architectures, is limited by computational demands and a lack of supporting tools. Here, we introduce *fibertools*, a state-of-the-art toolkit that features a semi-supervised convolutional neural network for fast and accurate identification of m6A-marked bases using PacBio single-molecule long-read sequencing, as well as the co-processing of long-read genetic and epigenetic data produced using either PacBio or Oxford Nanopore sequencing platforms. We demonstrate accurate DNA-m6A identification (>90% precision and recall) along >20 kilobase long DNA molecules with a ∼1,000-fold improvement in speed. In addition, we demonstrate that *fibertools* can readily integrate genetic and epigenetic data at single-molecule resolution, including the seamless conversion between molecular and reference coordinate systems, allowing for accurate genetic and epigenetic analyses of long-read data within structurally and somatically variable genomic regions.

## Introduction

Highly accurate long-read single-molecule DNA sequencing has revolutionized the comprehensive assembly of phased genetic architectures, enabling the first complete human genome assemblies (Wenger *et al*., 2019; Vollger *et al*., 2020; Nurk *et al*., 2022). In addition, long-read DNA sequencing natively identifies endogenously modified DNA bases, such as m6A and 5-methylcytosine (5mC), permitting the co-analysis of both genetic and DNA methylation features at single-molecule resolution (Marks *et al*., 2012; Clark *et al*., 2012; Murray *et al*., 2012; Loman *et al*., 2015; primrose: Predict 5mC in PacBio HiFi reads). Furthermore, using exogenous DNA methyltransferases to add DNA base modifications, such as in the context of single-molecule chromatin fiber sequencing (Stergachis *et al*., 2020; Lee *et al*., 2020; Abdulhay *et al*., 2020; Shipony *et al*., 2020; Altemose *et al*., 2022), permits the co-analysis of genetic, DNA methylation, and chromatin epigenetic features at single-molecule and single-nucleotide resolution.

Specifically, single-molecule chromatin fiber sequencing leverages non-specific methyltransferases to selectively stencil chromatin protein occupancy patterns directly onto their underlying DNA molecules in the form of modified bases. Modified bases along individual DNA molecules are then directly identified using PCR-free single-molecule sequencing. For example, during single-molecule, real-time (SMRT) sequencing, the identity of each base is determined by the fluorophore-labeled nucleotide that is incorporated as the polymerase replicates the base. In contrast, the modification status of each base is determined by signature changes in polymerase kinetics at and surrounding that base as it is replicated by the polymerase, such as elongation of the interpulse duration (IPD) due to polymerase pausing at modified bases (Flusberg *et al*., 2010) (**Fig. 1a, S1**). Recently developed tools leverage these polymerase kinetic parameters to identify 5mC within specific sequence contexts (Tse *et al*., 2021; primrose: Predict 5mC in PacBio HiFi reads), genomic positions with consistent m6A signal across multiple sequencing reads (Marks *et al*., 2012; Clark *et al*., 2012; Murray *et al*., 2012), total adenine methylation levels along very short sequencing reads (Kong *et al*., 2022), and m6A-modified bases at single-molecule resolution along only short ∼2 kilobase DNA molecules (ipdSummary, SAMOSA-ChAAT) (Abdulhay *et al*., 2023; Clark *et al*., 2012). However, accurately identifying DNA-m6A along multi-kilobase reads is largely unsolved, as existing approaches either have poor sensitivity/specificity (**Fig. S2**, **S3**), require excessive compute and storage resources (**Fig. S4**), and/or are reliant on input files (i.e., subreads) no longer available with modern sequencing chemistries.

**Figure 1.**
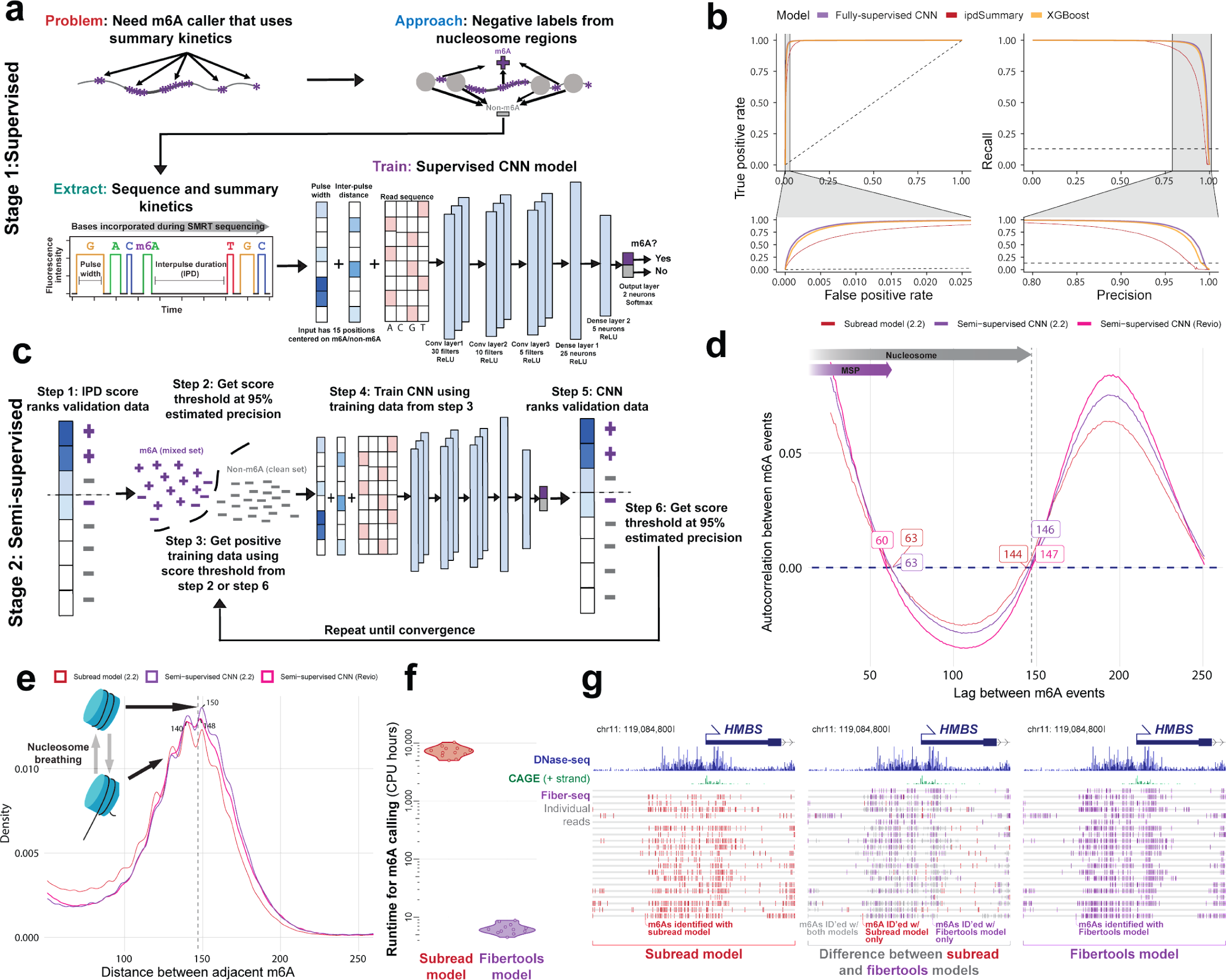
Accurate identification of m6A with supervised machine learning (ML) and refinement with semi-supervised ML. **a)** Methodology for generating training data and identifying m6A modifications using PacBio HiFi (see *Methods* for details). **b)** Receiver operating characteristic and precision-recall curves for the CNN (purple), XGBoost (orange), and ipdSummary (red) models. Dashed lines indicate the performance of a random classifier. **c)** Methodology for semi-supervised machine learning (see *Methods* for details). **d)** Autocorrelation between m6A calls made by the subread (red), semi-supervised (purple), and Revio (pink) models. **e)** Density of the distance between adjacent m6A on the same chromatin fiber (10,000 reads) for the same datasets and models as in d. **f)** CPU hours used by fibertools (purple) and subread-based GMM model (red) for individual SMRT cells. Fibertools was run with GPU acceleration (NVIDIA A40), which is unavailable for the GMM model. **g)** Visualization of m6A calls in the *HMBS* locus that are unique to fibertools (purple), unique to the subread GMM model (red), or shared by both (gray). Reads are sorted by the number of CCS passes. DNase-seq (ENCODE July 2012 Freeze) and CAGE signal is shown above.

Furthermore, although extensive tooling exists for processing short-read epigenetic data relative to reference coordinates (BEDOPS, BEDTools, etc.) (Neph *et al*., 2012; Quinlan, 2014), comparable tools for processing long-read epigenetic data are limited in their ability to leverage the rich epigenetic and genetic data embedded within long-read sequencing data (Razaghi *et al*., 2022). Specifically, tools for processing long-read epigenetic data need to operate in four dimensions: (1) they must process multiple types of genetic and epigenetic information present on a single read (e.g., DNA base, mCpG, DNA-m6A, inferred epigenetic marks, etc.); (2) they must capture this information across multiple reads mapping to a given reference position; (3) they must capture how this information co-occurs along each read in both reference and molecular coordinate systems; and (4) they must capture all of this information across the various haplotypes mapping to the same position within a reference.

Here, we introduce a semi-supervised machine learning approach for accurately identifying DNA-m6A in PacBio sequencing along multi-kilobase reads that permits the accurate learning of modified DNA bases from noisy training data - a common occurrence with single-molecule sequencing data owing to inherent biological heterogeneity in DNA methylation status between individual DNA molecules. Furthermore, we introduce a comprehensive toolkit for co-processing long-read genetic and epigenetic data designed for use across sequencing platforms.

## Results

Building an accurate tool for DNA-m6A identification requires a training dataset of multi-kilobase reads with both methylated and unmethylated adenines across diverse sequence contexts (i.e., all possible 7-mers containing a central adenine) and methylation density contexts (i.e., isolated or clustered DNA-m6As). Because creating such a dataset is not achievable using synthetic DNA or fully methylated and unmethylated samples, we leveraged DNA from single-molecule chromatin fiber sequencing reactions (i.e., Fiber-seq) as the basis for training. Specifically, Fiber-seq uses non-specific m6A-MTases to selectively mark sites of protein occupancy along individual DNA molecules via m6A-marked bases. Because protein occupancy is highly heterogeneous across chromatinized DNA (**Fig. S5**), each DNA molecule contains methylated adenines within diverse sequence and methylation density contexts (**Fig. S6**). Furthermore, we can employ chromatin features, such as nucleosome occupancy, to bolster training and validation, making Fiber-seq well suited for training a general-purpose DNA-m6A caller given labeled data.

To generate initial positive and negative labels, we used a previously published DNA-m6A caller (referred to here as the “subread model”) (Dubocanin *et al*., 2022). The subread model improves upon ipdSummary by using ipdSummary’s IPD normalization for sequence context (ipdRatios), followed by a Gaussian mixture model (GMM) to identify adenines with ipdRatios that significantly deviate from the expected distribution of unmethylated adenines (i.e., m6A-modified bases) (Dubocanin *et al*., 2022) (**Fig. S7**). These calls are then used to identify m6A-modified bases (positive labels). Negative labels are drawn from regions with extended stretches devoid of m6A corresponding to inferred nucleosome-occluded regions (**Fig. 1a**, *Methods*).

All existing DNA-m6A callers (i.e., ipdSummary, SAMOSA-ChAAT, and the “subread model”) require subreads. However, modern SMRT sequencing chemistries do not produce subreads and only produce summary kinetic information, making these existing tools largely obsolete. Consequently, we designed *fibertools* using a two-staged training approach. In stage 1 of training (**Fig. 1a,b**), we used a fully supervised training regime to validate that m6A calls can be generated using only summary kinetics, bypassing the requirement of all other PacBio m6A callers for individual subread kinetics. Using the dataset described above, we independently trained two machine-learning models, XGBoost (Chen and Guestrin, 2016) and a fully-supervised convolutional neural network (fully-supervised CNN), and evaluated their performance on a held-out dataset from a separate sequencing experiment (**Fig. 1a,b**, and *Methods*). Both the fully-supervised CNN and XGBoost models maintained high precision and recall, with average precision over 97% (**Table S1**). As a comparison, we benchmarked the models against ipdSummary (we excluded SAMOSA-ChAAT from benchmarking as it was only optimized for a single deprecated polymerase chemistry). Both our CNN and XGBoost models outperformed ipdSummary in terms of both AUPR and AUROC. At a 95% precision threshold, the fully-supervised CNN model has a recall of 89.6%, whereas the recall for ipdSummary is only 47.4%. Thus, the fully-supervised CNN nearly doubles the number of m6A identifications over ipdSummary at this precision threshold. Notably, the fully-supervised CNN and XGBoost models are ∼1,000x faster than the subread model used to generate the training dataset. Overall, the CNN model had the best performance, and we used it as the basis for stage 2 of training as well as subsequent improvements and validation.

In stage 2 of training, we sought to polish the architecture of our supervised CNN beyond the capabilities of existing tools and training datasets. To do this, we used a semi-supervised training regime (**Fig. 1c**) inspired by well-established methods in the field of proteomics (Käll *et al*., 2007; Fondrie and Noble, 2021) to overcome the limitations of deriving training data from existing models. The resulting semi-supervised model (referred to onwards as “*fibertools*”), allows for the possibility that the positive labels in the training dataset are incorrect (**Fig. 1c**, *Methods*). To evaluate this model, we established a series of biological validations to test the performance of the semi-supervised training since the direct use of labels from training data in semi-supervised machine learning does not provide an accurate assessment of performance.

First, we assessed the accuracy of *fibertools* for identifying nucleosome footprints along Fiber-seq data. Using single-molecule m6A calls with a predicted precision of >95% (*Methods*), we performed an autocorrelation analysis. Compared to the subread and fully supervised models, *fibertools* more accurately recapitulated the exact length of nucleosomes (147 bp) (**Fig. 1d**) (Luger *et al*., 1997) and showed an overall higher amplitude autocorrelation, consistent with higher quality identification of m6A with nucleosome patterning characteristic of human chromatin. In addition, comparing the distance between adjacent m6A methylation marks in human Fiber-seq data demonstrated clear oscillatory patterns suggestive of nucleosome breathing (Polach and Widom, 1995; Anderson and Widom, 2000; Hall *et al*., 2009), further indicative of high-quality m6A identification (**Fig. 1e**).

Second, we evaluated the false positive rate (FPR) of *fibertools* using whole-genome amplified (WGA) DNA that lacked m6A (**Fig. 2a**). Our findings indicated an FPR of 0.23% at the model-predicted precision level of >95% (**Fig. 2b, S8**). Notably, given this FPR, *fibertools* is not suited for identifying non-specific genomic m6A events within species with low-level endogenous m6A (Kong *et al*., 2022; Debo *et al*., 2023).

**Figure 2.**
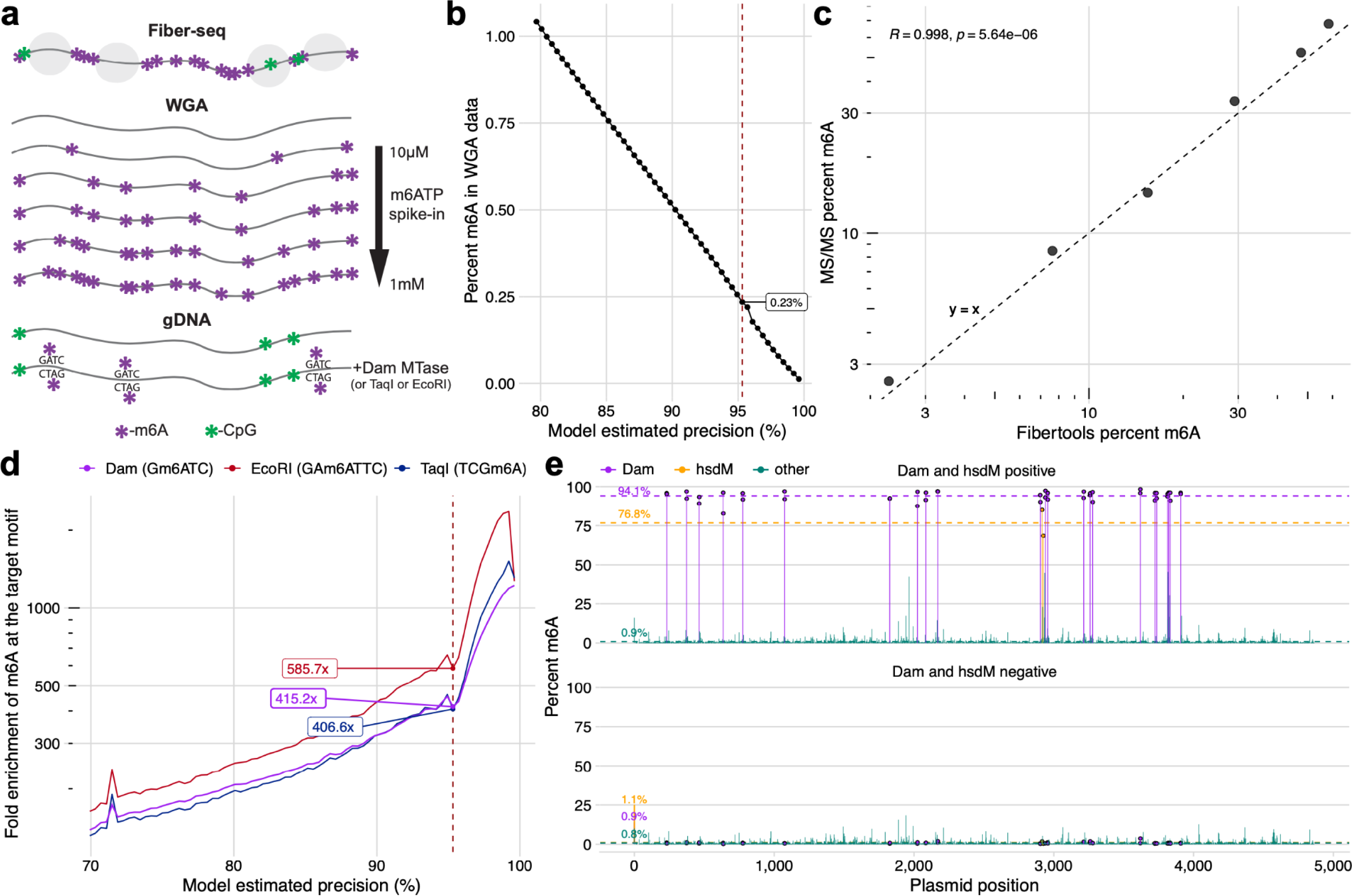
Biological validation of the semi-supervised m6A caller. **a)** Description of biological samples used for validation of fibertools. **b)** Percent of methylated adenines called by fibertools relative to all adenines in a whole-genome amplified (WGA) negative control as a function of the estimated precision reported by fibertools. This serves as an estimate of the false positive rate. The red line marks the default threshold used by fibertools. **c)** Percent m6A as determined by UHPLC–MS/MS (y) and fibertools (x) at the default precision level for WGA samples with varying levels of m6ATP spiked-in. The text (upper left) indicates the value of the Pearson’s correlation coefficient and the P value from a two-sided t-test without adjustment for multiple comparisons. **d)** Enrichment of m6A calls within targeted motifs of three motif-specific methyltransferases [Dam (blue), EcoRI (orange), and TaqI (green)] as a function of fibertools estimated precision. **e)** Methylation percent at recognition sites for *Dam* (purple), *HsdM* (orange), and other sites (green) among all sequencing reads of a plasmid grown in a dam^+^/hsdM^+^ bacterial strain (top) compared to a dam^-^/hsdM^-^ negative control (bottom). Dotted lines show the average across each category.

Third, we evaluated the ability of *fibertools* to accurately quantify the total amount of m6A within a sample. Specifically, we spiked varying levels of m^6^ATP into a WGA reaction (**Fig. 2a**) and employed ultra-high-performance liquid chromatography tandem mass spectrometry (UHPLC–MS/MS) to determine the percentage of methylated adenines with respect to all adenines (Methods). We then sequenced these samples, applied *fibertools*, and found a strong correlation (Pearson=0.998, p-value=5.2e-6) between our method and mass spectrometry (**Fig. 2c**).

Fourth, since the above validations demonstrate that m6A calls from *fibertools* recapitulate bulk chromatin features and total m6A content measured by UHPLC–MS/MS, we next evaluated the precision of *fibertools* for identifying isolated m6A events, which is relevant for m6A calls within small internucleosomal regions. Using *fibertools* to predict m6A on genomic DNA treated with motif-specific methyltransferases, we found that m6A calls were enriched by 415-fold, 586-fold, and 407-fold in motifs specific to Dam, EcoRI, and TaqI, respectively (**Fig. 2d**), consistent with precise m6A calls. The ability to call single m6A events is biologically relevant for fiber-seq as 10-25% of all internucleosomal linker regions (i.e., methyltransferase-sensitive patches, MSPs) within a Fiber-seq dataset contain only a single m6A separating adjacent nucleosomes (**Fig. S9**).

Fifth, the precision of *fibertools* in identifying single m6A events indicated it might be useful for identifying endogenously m6A-modified bases within bacteria, a context that also provides a true positive validation set as nearly 100% m6A methylation can be expected within methyltransferase-specific motifs. Notably, DNA isolated from bacteria expressing both the Dam and HsdM methyltransferases exhibited m6A at 94.1% of the target Dam sites for these methyltransferases, indicating a false negative rate of less than 6% in this sequence context (**Fig. 2e**). In contrast, DNA from bacteria lacking these methyltransferases (Anton et al. 2015) exhibited m6A at <1% of adenines, consistent with our prior FPR estimate.

Sixth, we evaluated the accuracy of *fibertools* for quantifying m6A-marked chromatin architectures along multi-kilobase reads. Current SMRT cell chemistries target sequencing reads ∼20 kb in length, yet traditional subread-based m6A models were designed for reads of only ∼2kb in length. We demonstrate that in comparison to existing subread-based models, *fibertools* substantially reduce false-negative methylation calls along multi-kilobase reads (**Fig. 3, S10**), enabling the accurate quantification of m6A-marked chromatin architectures along reads >25 kb in length (**Fig. 3b**).

**Figure 3.**
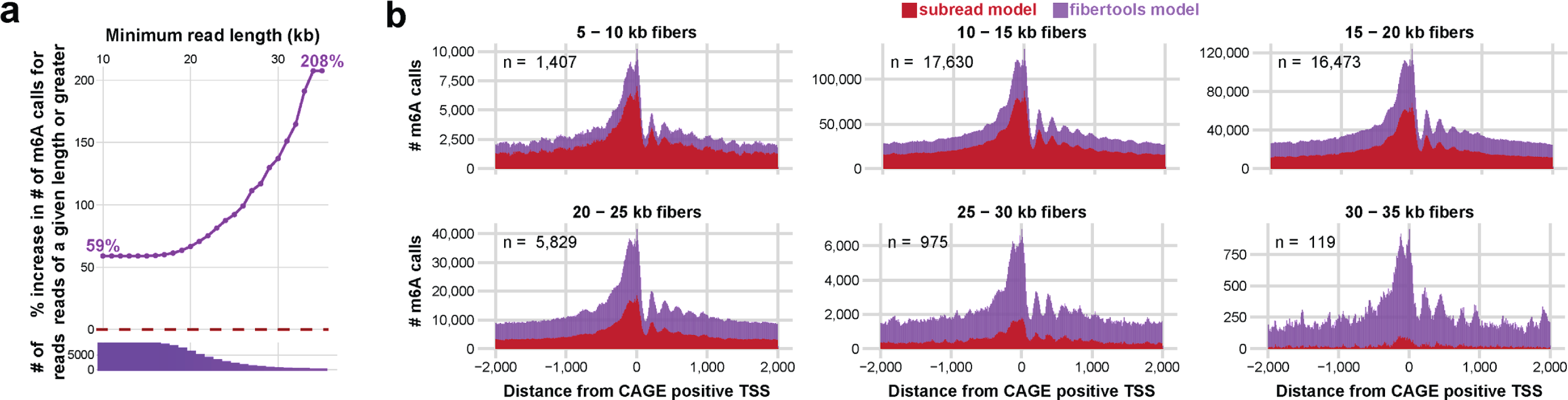
Increased m6A calling on long reads (>20kb) via fibertools. **a)** Percent increase in fibertools m6A calls over the GMM model as a function of the minimum read length of the underlying sequencing data. The histogram below shows how many reads were used to calculate each percent increase. **b)** Comparison of fibertools and the subread model for m6A calling over CAGE-positive TSS in K562 cells across the genome, separated by read length. Reads are matched between fibertools and the subread model, and the number of Fiber-seq reads used in the calculation of each size range (n) is indicated.

Collectively, these biological validations provide strong evidence that *fibertools* is highly accurate and specific in identifying m6A events using PacBio HiFi data. Importantly, the semi-supervised training design of *fibertools* enables it to readily adapt to new sequencing chemistries, which often contain updated polymerases that may differ in their kinetic values (**Fig. S1**). For example, we used calls from the model for the PacBio Sequel II 2.2 chemistry as the initialization point for training a semi-supervised model for the PacBio Revio chemistry - demonstrating that this new Revio model is similarly highly accurate in identifying m6A events (**Fig. 1d,e**).

Having established that *fibertools* can identify highly accurate m6A events using PacBio HiFi data, we next sought to extend *fibertools* to enable the simultaneous processing of genetic, cytosine methylation, and adenine methylation data. To accomplish this, we designed *fibertools* to integrate m6A calls directly into the BAM format using the MM and ML tags (**Fig. S11**). Next, we optimized *fibertools* using a compiled language, which we provide as a single binary (*ft*) accessible through bioconda (package “*fibertools-rs*”). Of note, *fibertools* can process individual Revio SMRT cells in 15-24 CPU hours and Sequel II SMRT cells in 5-8 CPU hours, a >1,000-fold increase in speed compared to the previous pipeline when using GPU acceleration (>150-fold increase without GPU) (**Fig. 1f, Table S2**).

We next extended the utility of *fibertools* to perform fundamental operations necessary for processing single-molecule epigenetic and genetic data produced using either PacBio and ONT sequencing platforms (i.e., *fibertools add-nucleosome*, *fibertools extract,* and *fibertools center*).

*Fibertools add-nucleosome* enables the identification of stretches of unmethylated adenine bases, which are stored directly in the BAM file using custom flags. It processes 10 million ∼20kb reads in just 4.6 CPU hours. Importantly, *fibertools add-nucleosome* works seamlessly with Fiber-seq data sequenced using either a PacBio or ONT instrument. For example, the application of *fibertools add-nucleosome* to a Fiber-seq library sequenced on an ONT R10.4 flow cell and base called using Dorado v0.4.2 enabled the robust identification of clear nucleosomal patterns, despite the substantial decrease in m6A and DNA base identification accuracy with ONT sequencing (**Fig. 4**).

**Figure 4.**
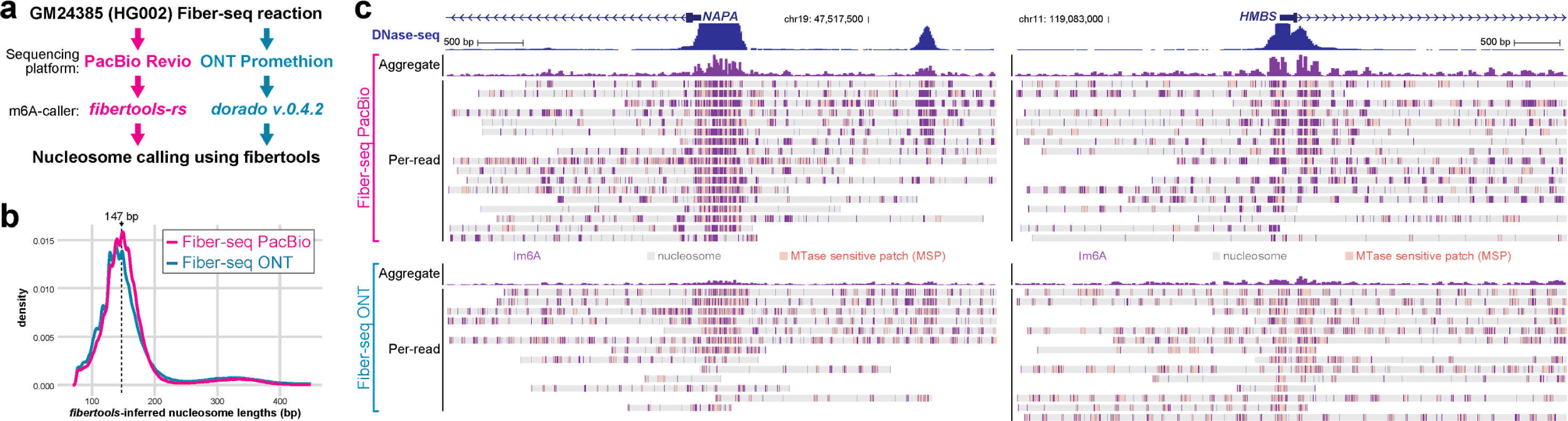
Fibertools nucleosome calling with PacBio and ONT Fiber-seq. **a)** Data processing pipelines for fibertools nucleosome calling with *fibertools*. **b)** Density of nucleosome lengths called by *fibertools* for PacBio (pink) and ONT (blue) Fiber-seq (n = 500,000 nucleosomes). **c)** Visualization of the *NAPA* and *HMBS* representative loci for PacBio (top) and ONT (bottom) Fiber-seq. m6A calls from *fibertools* (PacBio) or *Dorado* (ONT) are represented by vertical purple dashes, along with nucleosome (gray) and MTase-sensitive patch (MSP) (orange) calls from *fibertools*.

*Fibertools extract* enables multithreaded conversion of genetic and epigenetic BAM features into plain text formats regardless of upstream tooling with coordinates in either reference or molecular space. For example, nucleosomes, m6A, and 5mC methylation can all be extracted into BED12 format to visualize with the UCSC genome browser. Alternatively, all of these features and more can be extracted into a unified table for custom downstream analysis or visualization.

*Fibertools center* enables the processing of single-molecule genetic and epigenetic data (i.e., DNA base, mCpG, DNA-m6A, inferred epigenetic marks, etc.) relative to a set of reference genomic coordinates while maintaining how these features co-occur along each read using both reference and molecular coordinate systems - addressing a need that is unique to long-read epigenetic studies. To demonstrate the utility of *fibertools center,* we applied it to address two fundamental biological questions that require the integration of long-read epigenetic and genetic data: (1) the co-occupancy of transcription factor (TF) binding elements along individual DNA molecules (**Fig. 5**); and (2) the relationship between somatic DNA variability and overlying altered epigenetic architecture along individual DNA molecules (**Fig. 6**).

**Figure 5.**
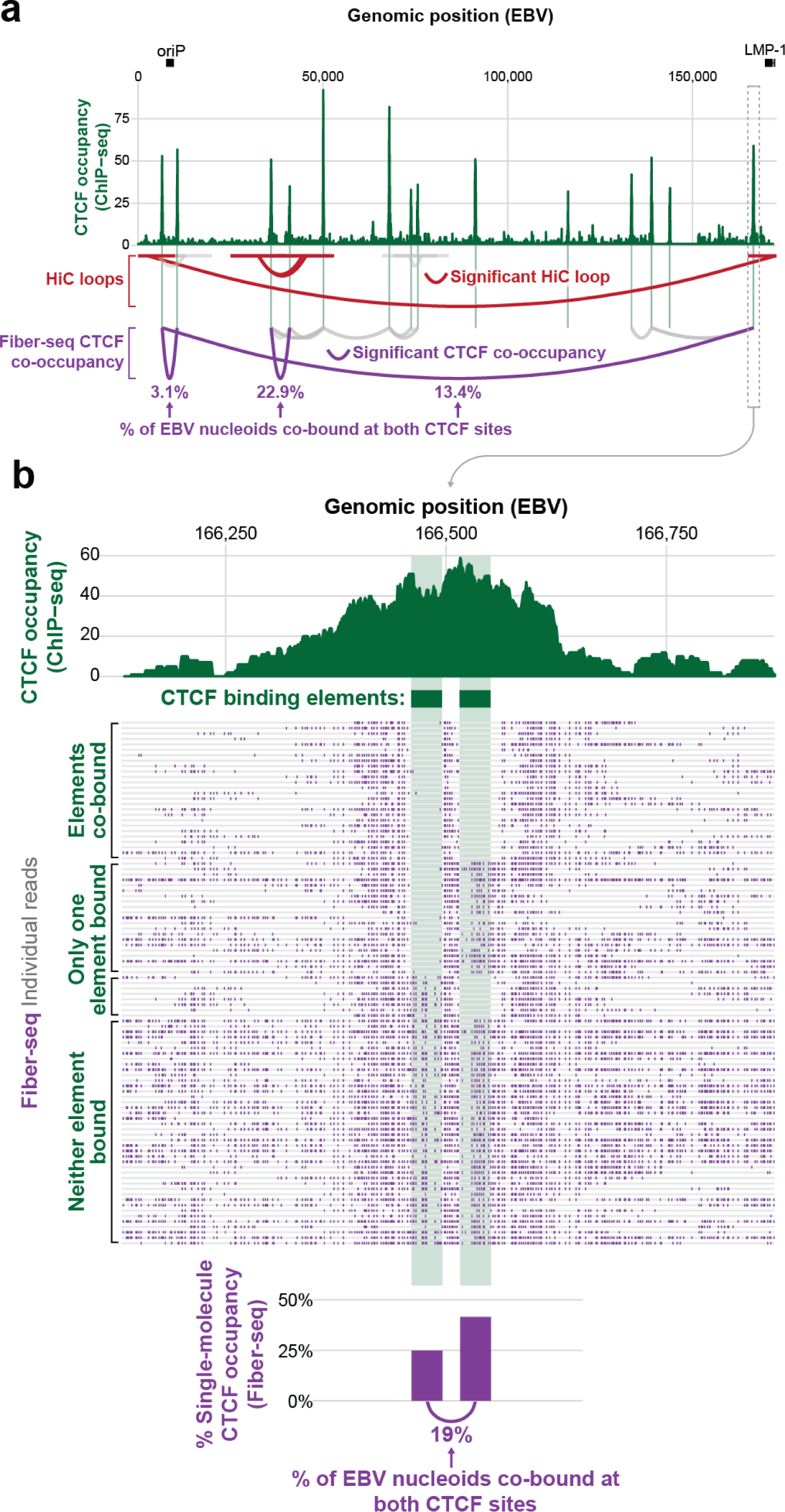
CTCF co-occupancy along the EBV genome. **a)** CTCF ChIP-seq (green), significant (red) or insignificant (gray) HiC loops (Morgan *et al*., 2022), and significant CTCF site co-occupancy by Fiber-seq (purple) along the EBV genome. The significance of CTCF site co-occupancy was determined by comparing the expected number of co-occupied fibers to the observed number using Fisher’s Exact test (see **Table S6** for exact counts and p-values). **b)** Zoom-in of the indicated CTCF peak, which contains two CTCF binding elements. Single-molecule occupancy and co-occupancy from Fiber-seq are shown below.

**Figure 6.**
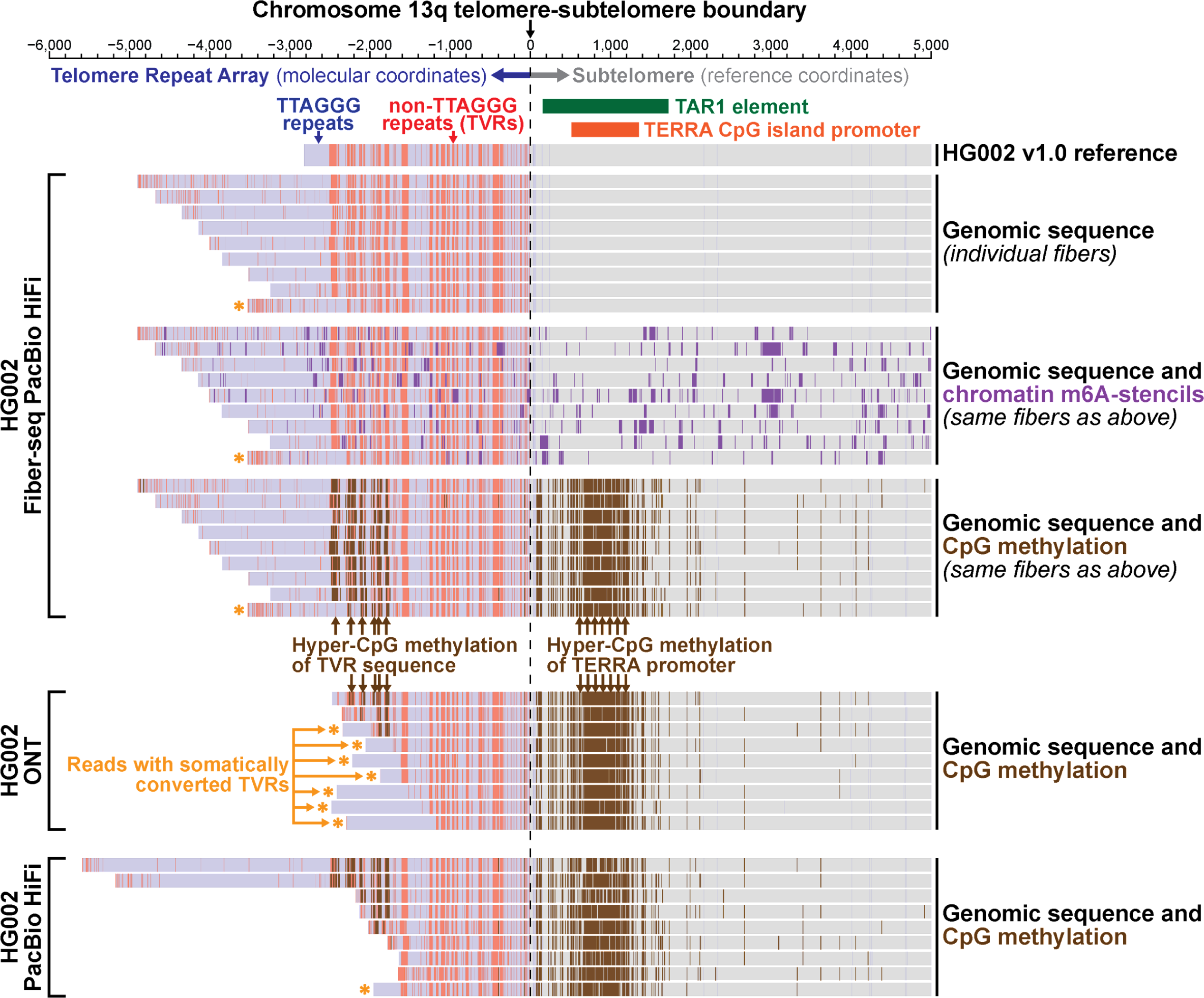
Organization of the HG002 telomere. **Top**) HG002 PacBio Fiber-seq with genetic variants, m6A methylation (purple), and CpG methylation (brown) overlaid for fibers overlapping the telomere of chromosome 13q. Telomeric sequences (blue), telomeric variants (red), and non-telomeric sequences (gray) are highlighted to show telomeric genetic variation. (**Middle**) ONT and (**bottom**) PacBio standard sequencing of HG002 telomeres (Zook *et al*., 2020) with CpG methylation overlaid on chromatin architecture.

First, we applied Fiber-seq and *fibertools center* to resolve the role of CTCF co-occupancy in guiding higher-order chromatin architectures along the ∼175 kbp Epstein-Barr virus (EBV) genome. CTCF elements within the EBV OriP and *LMP* loci are known to form a cohesin-dependent loop important for maintaining viral latent cycle gene expression (Arvey *et al*., 2012; Morgan *et al*., 2022). Using *fibertools,* CTCF footprints were resolved on individual DNA molecules (Yin *et al*., 2017), and we determined that the CTCF ChIP-seq peak within the EBV *LMP* locus is predominantly mediated by CTCF co-occupying two immediately adjacent CTCF binding elements along the same DNA molecule (**Fig. 5**). In addition, we observed that EBV nucleoids bound by CTCF within the *LMP* locus are also preferentially co-bound by a single CTCF element within the EBV OriP locus, which is located ∼12 kbp away (**Fig. 5a**). However, only 13.4% of EBV nucleoids are co-bound by CTCF at both the OriP and *LMP* loci (**Fig. 5b**), setting an upper limit on the proportion of EBV nucleoids that are directly structured by CTCF-mediated three-dimensional looping between these two loci in a stable manner. Notably, dynamic CTCF-mediated looping has been observed along the nuclear genome (Gabriele *et al*., 2022), and it is possible that these CTCF co-bound EBV nucleoids may represent similarly dynamic CTCF-mediated looping along the EBV genome.

Second, we applied *fibertools center* to resolve the relationship between somatic DNA variability and overlying altered epigenetic architecture along telomeric and sub-telomeric regions using Fiber-seq PacBio HiFi, standard PacBio HiFi, and standard ONT sequencing data. Telomere repeat arrays exhibit substantial per-molecule somatic alterations in both their length and sequence content (Dubocanin *et al*., 2022) and are known to be transcribed into Telomeric Repeat-containing RNA (TERRA) via a CpG island promoter located within a TAR1 repeat positioned adjacent to the majority of telomere repeats (Azzalin *et al*., 2007). The substantial molecule-to-molecule heterogeneity in the DNA content of individual telomere repeats originating from the same chromosome arm requires the use of a molecular coordinate system for their appropriate analysis (i.e., the original positions from the sequenced read, without adjustments for alignment to the reference sequence). Application of *fibertools center* to Fiber-seq data from HG002 cells aligned to the HG002 reference genome enabled the resolution of both genetic and epigenetic architectures of sub-telomeric regions in reference coordinates and telomere repeat arrays in molecular coordinates by using the telomere-subtelomere boundary as the centering point. Importantly, by using molecular coordinates to display the telomere repeat array, we were able to identify multiple non-TTAGGG telomere variant repeats (TVRs) (Baird *et al*., 1995; Allshire *et al*., 1989) absent from the HG002 reference sequence (**Fig S12**). Overall, this analysis exposed that the sub-telomeric TAR1 element was hyper-CpG methylated and lacked chromatin accessibility within HG002 cells (**Fig. 6**), in contrast to CHM13 cells (Dubocanin *et al*., 2022). Notably, this region of hyper-CpG methylation extended into TVRs harboring CpG dinucleotides within the telomere repeat array itself (Baird *et al*., 1995; Allshire *et al*., 1989), which we validated by applying *fibertools* to additional genomic ONT and PacBio HiFi sequencing data from HG002 (Zook *et al*., 2020). TVRs can undergo somatic conversion back into TTAGGG repeats, likely via somatic shortening and subsequent telomerase-mediated elongation (Dubocanin *et al*., 2022). Integrating the genetic and epigenetic data along individual telomere fibers identified numerous fibers that had undergone such somatic conversions, resulting in the loss of CpG methylation within that region of the telomere repeat array. However, hyper-CpG methylation of the TAR1 element and the rest of the telomere repeat array was maintained on these fibers, suggesting that CpG methylation of TVRs may be a bystander effect of TAR1 hyper-CpG methylation within HG002 cells.

## Discussion

In summary, *fibertools* provides foundational tooling for processing long-read genetic and epigenetic data. Specifically, *fibertools* enables highly accurate single-molecule DNA-m6A identification using PacBio sequencing with a 1,000-fold improvement in speed (**Fig. 1f**), enabling highly accurate single-molecule chromatin fiber sequencing and endogenous bacterial adenine methylation identification (**Fig. 1d, 2e, S13**). In addition, *fibertools* enables the synchronous processing of multiple types of genetic and epigenetic information present within BAM files produced using either PacBio or ONT sequencing platforms at single-molecule and single-nucleotide resolution (**Fig. 4**). Furthermore, *fibertools* enables the seamless conversion between molecular and reference coordinate systems, allowing for accurate genetic and epigenetic analyses of long-read data within structurally and somatically variable genomic regions (**Fig. 6)**. Finally, fibertools is written in a compiled language and is available as a single binary accessible through bioconda (package “fibertools-rs”) and PyPI (package “pyft”), enabling its broad utility within existing computational pipelines for processing long-read data. Recent advances in the cost, accuracy, and throughput of long-read sequencing have enabled the broader adoption of this technology across the genetics community, and we anticipate that *fibertools* will serve as an integral tool for processing long-read sequencing data.

Finally, the semi-supervised training structure that *fibertools* introduces for identifying m6A-marked bases will likely prove useful for the accurate identification of other base modifications using long-read sequencing (**Fig. 1c**, *Methods*). Specifically, this approach accounts for imperfect training data, which is a core feature of long-read sequencing data owing to the inherent biological per-molecule heterogeneity in the distribution of modified bases. As such, we anticipate that this approach can both improve the accuracy of identifying core base modifications in humans, as well as novel base modifications in bacteria. Furthermore, we demonstrate that models trained using a semi-supervised structure can readily adapt to updated sequencing chemistries, making this training approach highly advantageous for long-read sequencing chemistries, which are undergoing frequent iterative development cycles (**Fig. 1d,e**).

In summary, we present *fibertools* as a toolkit to facilitate the identification and analysis of long-read genetic and epigenetic sequencing data and establish semi-supervised machine learning as an adaptable tool for accurately identifying base modifications using long-read sequencing.

## Methods

### Initial m6A calling with ipdSummary followed by filtering with a GMM

To initially identify m6A, we use a previously published protocol (Dubocanin *et al*., 2022) with the following details and modifications. Raw PacBio subread BAM files were converted into CCS (circular consensus sequence) reads using pbccs (v6.0.0) (https://ccs.how/) with average kinetics information included. Then subreads were aligned to their respective CCS reads using actc (https://github.com/PacificBiosciences/actc). The resulting alignments were passed to ipdSummary (v3.0) (https://github.com/PacificBiosciences/kineticsTools/) to identify positions with m6A modifications with the CCS-specific extracted genomic sequence as the reference and these additional flags: --pvalue 0.001 --identify m6A. For each read, we then trained a two-component Gaussian mixture model (GMM) from the ipdRatios generated by ipdSummary for all adenine bases within the read (push_m6a_to_bam.py). Adenine bases were then classified as m6A modified if the probability that the ipdRatio came from the larger distribution was greater than or equal to 0.99999999. All code to repeat these steps is made available as a single snakemake (Köster and Rahmann, 2012) pipeline on GitHub (https://github.com/StergachisLab/fiberseq-smk).

### Hidden Markov model nucleosome calling

To identify nucleosomes initially, we use a previously published protocol (Dubocanin *et al*., 2022) with the following details and modifications. We used m6A calls to train a two-state (nucleosome, non-nucleosome) Hidden Markov model (HMM) with a standardized post-hoc correction. The input data for the HMM skipped all G/C bases and considered only the A/T sequence so that GC-rich regions without adenine would not falsely be called nucleosomes. The HMM was then trained using the Baum-Welch algorithm (Baum *et al*., 1970) using a subsample of 5,000 CCS reads per SMRT cell, and the trained model was subsequently applied across all fibers from that SMRT cell using the Python package pomegranate (Schreiber, 2017). In tandem with the HMM delineation of nucleosomes, we also applied a simplified approach that looked for unmethylated stretches larger than 85 bp in length (irrespective of A/T content). We used these ‘simple’ calls to refine our HMM nucleosome calls by splitting large HMM nucleosomes that contained multiple ‘simple’ calls. We also refined the terminal boundaries of each nucleosome by bookending it to the nearest m6A call within 10 bp, if present. Nucleosome calling is available as part of a snakemake pipeline (https://github.com/StergachisLab/fiberseq-smk) and encodes the resulting calls in the custom BAM flags ns (nucleosome start) and nl (nucleosome length), which can be extracted using fibertools-rs.

### Generating positive and negative labels for training and validation data

To generate positive labels, we used the previously existing calls made using the GMM-filtered calls from ipdSummary in our snakemake pipeline. Negative labels were drawn from nucleosome regions as defined by the HMM since we had increased confidence that these regions should be inaccessible due to the presence of a nucleosome. For each SMRT cell, we selected ∼350,000 CCS reads and randomly sampled 5% of available positive and negative m6A positions within each read to reduce the number of adjacent and, therefore, non-independent calls in our training data. This resulted in a dataset of ∼100 million negative labels and ∼8 million positive labels for each SMRT cell. We note that by selecting negative labels from within nucleosomes, we allow for the possibility for our models to outperform ipdSummary and the GMM correction even though we draw positive labels from these tools. For each label in our dataset, we included a 15 bp window centered around the adenine encoding the sequencing information with one-hot encoding and the CCS kinetics information across the same 15 bases for pulse width and interpulse duration resulting in a 6 × 15 matrix for each labeled position. The 15 bp window and inclusion of pulse width and interpulse duration were selected based on the ablation study of the supervised model (**Fig. S14**). Code to generate training data from CCS reads is available on GitHub (https://github.com/mrvollger/m6A-calling).

### Training, validation, and testing datasets for machine learning

For the 2.2 PacBio sequencing chemistry, we established three separate datasets, each from a different sequencing run, for developing our models. The three datasets were used for training, validation, and a completely held-out testing dataset for determining final accuracies. Data for v2.2 chemistry was generated from previously described samples (K562, CHM1) treated with 200 units of Hia5 for 10 minutes at 25°C (Dubocanin *et al*., 2022). Data for v3.2 chemistry was generated from a K562 Fiber-seq sample (described below), and Revio data was provided by PacBio.

### Architectures and training of machine-learning models (XGBoost, CNN, and semi-supervised CNN)

We trained an XGBoost model with a binary logistic objective function using our training and validation datasets. We selected the following hyperparameters using threefold cross-validation; learning rate of 1 (gamma), maximum tree depth of 8, minimum child weight of 100, and 150 estimators. The code to repeat this training is available on GitHub (https://github.com/fiberseq/train-m6A-calling).

### Convolutional neural network

We trained a CNN to predict m6A in Fiber-seq HiFi reads. Input to the CNN model is the 6×15 matrix described above. The model has three convolutional layers with 30, 10 and 5 filters of size 5, 5, and 3, respectively. A dense layer of size 25 × 5 and an output layer of size 5 × 2 follow the convolutional layers. All internal layers have ReLU activation, and the output layer has softmax activation. The output layer has two classes, one for m6A and one for unmethylated adenines. Each class generates scores between 0 and 1 for each input matrix, where scores close to 1 denote high confidence that the input belongs to that class. For example, unmethylated adenines score close to 0 for the m6A class, and methylated adenines score close to 1. The CNN optimizes a binary cross-entropy loss function using the Adam optimizer (Kingma and Ba, 2014). We trained the model iteratively for 30 epochs, where the training data was input in random batches of 32 in each epoch. See **Table S3** for the size of training and validation datasets from different Fiber-seq chemistries for supervised training. The code to repeat this training is available on GitHub (https://github.com/fiberseq/train-m6A-calling).

### Semi-supervised convolutional neural network

We derive m6A labels from GMM-filtered ipdRatios from ipdSummary. These labels can contain false positives. The lack of clean m6A labels makes the supervised training approach less suitable since it assumes that accurate labels are available. Therefore, we developed a semi-supervised approach, which assumes that our m6A class has a mixed population of true and false positives and our non-m6A class is a clean set. Our training approach is derived from the Percolator and Mokapot proteomics tools for identifying peptides from tandem mass spectrometry data (Käll *et al*., 2007; Fondrie and Noble, 2021). This approach yields a classifier with m6A calls at a target precision. The semi-supervised algorithm is outlined in *Supplementary Methods* and *Supplementary Algorithm 1*. First, we split our dataset into training and validation sets stratified by class labels (see **Table S3** for the size of training and validation datasets from different Fiber-seq experiment chemistries for semi-supervised training). Then our method proceeds in two phases. In the first phase, we use the IPD score of the central base as a classifier and generate an m6A classification score for all examples in the validation set. The classification score ranks the validation examples, and precision is computed at every score threshold. At the end of this phase, we select a score threshold to achieve the target precision. In this work, we use a target precision of 95%. The second phase is iterative, and each iteration consists of three steps. The first step is selecting a high-confidence m6A training set using the current score threshold. The second step consists of training a CNN model on this training data. In the final step, the validation data is rescored using the trained CNN model from the second step, and a new score threshold is generated at 95% precision with the rescored validation data. In the case of a successful second phase training, the number of positives in the validation data identified at target precision increases with every iteration and plateaus when most m6A examples in the validation data have been identified. We define two conditions for convergence, both of which must be satisfied. First, more than 70% of putative m6A calls from the validation set have been identified. Second, the number of additional m6A calls in a new iteration is less than 1% of the total putative m6A calls. In practice, it took 12, 11, and 3 iterations of phase two training to converge 2.2, 3.2, and Revio chemistry Fiber-seq experiments, respectively (**Fig. S15**). The code to repeat this training is available on GitHub (https://github.com/fiberseq/train-m6A-calling).

### Encoding and selecting precision levels for m6A calling

Using the approach outlined in the semi-supervised method, we calculated the empirical precision using the validation data for every score output from the CNN model (see *Supplementary Methods* and **Table S4** for details). These precisions are then multiplied by 256 and rounded into an 8-bit integer following the BAM specification for the ML tag (https://samtools.github.io/hts-specs/SAMtags.pdf, (Li *et al*., 2009). We then chose the first value (244) with a precision greater than 95% (244/256) as the threshold for calling positive m6A events.

### Base calling for Oxford Nanopore

HG002 DNA prepared with the standard fiber-seq protocol (below) was sequenced on a PromethION flow cell (R 10.4.1) with a 5kHz sampling rate. Alignment to the HG002 or GRCh38 genome and DNA base and base modifications calling were performed with Dorado v.0.4.2 with the basecalling model dna_r10.4.1_e8.2_400bps_sup@v4.2.0, 5mC model V2, and 6mA model V3.

### Chemicals and materials used

1 M Tris-HCl (pH8, molecular biology grade ultrapure, ThermoScientific, cat#J22638-K2), 2-deoxyadenosine (A) (>99%, FisherScientific, cat#AAJ6388606), Benzonase (Millipore Sigma, cat#E1014), CpG methyltransferase M.SssI(NEB, cat#M0226), dNTP set (Neta Scientific, cat#GHC-28-4065-51), HMW DNA Extraction kit (Promega, Cat#A2920), M. SssI (NEB, cat#M0226S), N6-methyl-2-dATP (TriLink, cat#N-2025), N6-methyl-2-deoxyadenine (m6A) (>99%, FisherScientific, cat#AAJ64961MD), Nanosep (MWCO 3 kDa, Pall, cat#OD003C33), phosphodiesterase I (Worthington, cat#LS003926), Quick CIP (NEB, cat#M0525S), Ultrapure distilled water (Invitrogen, cat#10977015). Taq methyltransferase (NEB, cat#M0219S), Dam methyltransferase (NEB, cat#M0222S), EcoRI methyltransferase (NEB, M0211S), ProNex® Size-Selective Chemistry (Promega, cat#NG2001), PacBio Elution buffer (PacBio #101-633-500), ZORBAX Eclipse Plus C18 (Agilent, cat#959757-902). All reagents used for mass spectrometer analysis are molecular grade level or above. g-TUBEs, SMRTbell® prep kit 3.0, REPLI-g Mini kit (Cat# 150023).

### Preparation of calcium-competent ER2796 E. coli

A 10mL culture of the ER2796 E. coli strain (obtained from NEB) was grown overnight without antibiotics. 1mL of the overnight culture was added to 99mL of fresh LB media without antibiotics and incubated with shaking at 37°C and 200 rpm for 3-4 hours until OD reached 0.4. The culture was separated into two 50mL falcon tubes and placed on ice for 20 minutes before being pelleted by centrifugation at 4°C and 4000 rpm for 10 minutes. The supernatant was discarded, and the cell pellets were resuspended with 20mL of ice-cold 0.1 M CaCl2 and incubated on ice for 30 minutes. The cells were then pelleted again by centrifugation and the supernatant was discarded. The pellets were then combined by resuspending in 5mL of ice-cold 0.1M CaCl2 with 15% glycerol. Cells were aliquoted in 50ul aliquots, frozen in liquid N2, and stored at −80C.

### Plasmid DNA preparation

A pCS2+ plasmid containing flag-tagged mgfp5 cloned into the EcoRI/XhoI sites (a gift from Lea Starita) was transformed into two strains of competent E. coli, ER2796 (from NEB, see “Preparation of calcium competent ER2796 E. coli”) and NEB 5-alpha (NEB cat. no. C2987I), using a standard heat shock method. 50 µL of the chemically competent cells were thawed on ice and mixed with 50 ng of plasmid DNA. The mixture was placed on ice for 30 minutes before being heat shocked at 42°C for 30 seconds. After heat shock, the cells were immediately placed back on ice for 5 minutes. Following incubation, 950 µL of room temperature SOC media (NEB cat. no. B9020S) was added, and the cells were allowed to outgrow for 1 hour in the absence of selection. 50 µL of the outgrown cells were diluted in 5 mL of selective media and grown overnight with shaking at 37°C and 220 rpm. Specifically, the NEB 5-alpha cells were grown in LB media with 100 μg/mL ampicillin, while the ER2796 cells were grown in LB media with 50 μg/mL kanamycin + 100 μg/mL ampicillin. The following day, plasmid DNA was extracted from the bacterial cells using the Monarch Plasmid Miniprep Kit (NEB cat. no. T1010L) following the manufacturer’s protocol. The elution of plasmid DNA was done with sterile water and the concentration was measured using the Qubit 1X dsDNA HS Assay Kit (Invitrogen cat. no. Q33231).

### K562 cell culture

K562 cells were maintained in suspension in IMDM media supplemented with 10% FBS (HyClone, cat. no.SH30396.03IH25-40) and antibiotic (100 I.U/mL penicillin, 100ug/mL streptomycin, Gibco, cat. no. 15140122) at 37°C and 5% CO2 in T-75 flasks. Cells were split 1:10 every 3-4 days.

### gDNA preparation

Whole-genome DNA was extracted from K562 cells using HMW DNA extraction kit (Promega, Cat# A2920). gDNA was used as a template for all WGA samples. All DNA samples were rehydrated with 10 mM Tris-HCl, pH 8 (ThermoScientific, cat# J22638.K2) or eluted with ultrapure water unless specified.

### WGA preparation

WGA was performed with REPLI-G Mini kit (Cat# 150023) according to the manufacturer’s protocol. 0, 10, 25, 64, 160, 400, or 1000 µM of N6-methyl-2-dATP (TriLink, cat#N-2025) (m6dATP) final concentration was spiked into 50 µL of each WGA to generate samples with 0, 2.56, 8.46, 14.45, 33.42, 52.27, and 67.94% m6A samples, as calculated using mass spec according to an eleven point standard quantified every time with the samples during the same run (see below). Samples were purified with ProNex® Size-Selective Chemistry (Promega, cat#NG2001) by adding a 1:1.3 ratio by volume of sample to magnetic beads. Following incubation for 15 minutes, the sample was placed on a magnetic rack for three minutes, washed twice with 80% ethanol, and eluted in 50 µL of PacBio Elution buffer (PacBio #101-633-500). All m6dATP-spiked samples were purified three times with bead purification, and residual free m6dATP quantified by MS of the 10 and 160 µM m6dATP samples treated with Quick CIP only was near background levels.

### gDNA methylation with sequence-specific methyltransferases

Samples labeled with the sequence-specific adenine methyltransferases EcoRI (GAATTC) (NEB, M0211S), Dam (GATC) (NEB, cat#M0222S), or TaqI (TCGA) (NEB, cat#M0219S) were generated as follows: 400 ng of purified gDNA/purified WGA DNA was treated for one hour with 40U of EcoRI (37°C) (gDNA only), 10U of TaqI (65°C), or 8U of Dam (37°C) in supplied buffer and in the presence of 160 μM SAM. All samples were purified once by ProNex® Size-Selective Chemistry as described above but with a 1:0.8 ratio by volume of sample to magnetic beads.

### Fiber-seq

Non-specific methyltransferase, Hia5, was purified, and the activity was quantified as previously described (Stergachis *et al*., 2020). Fiber-seq samples were prepared as previously described (Stergachis *et al*., 2020).

### Library preparation

Samples were sheared in g-TUBEs (Covaris, cat# 520079) for four passes at 3200 RPM for 2-4 minutes in an Eppendorf 5424R centrifuge. Post-shear samples were quantified by Qubit dsDNA high-sensitivity assay (Qubit, cat# Q32851) following the manufacturer’s protocol. Multiplied library preparation was performed using the SMRTbell prep kit 3.0 (PacBio, cat# 102-141-700) and SMRTbell barcoded adapter plate 3.0 (bc2001-bc2019) (PacBio, cat# 102-009-200) according to the manufacturer’s instructions but with the following modifications: after barcoded adapter ligation, samples were incubated at 65°C for 10 minutes to heat-inactivate the ligase. Barcoded samples were pooled and purified with ProNex® Size-Selective Chemistry as described above, with a 1:1 ratio by volume of sample to magnetic beads. Following nuclease treatment, the library was purified first with a 1:3.1 ratio of sample to 35% v/v Ampure PB beads (PacBio, cat# 100-265-900)/PacBio elution buffer. A second purification was performed with a 1:1 ratio of sample to ProNex® Size-Selective Chemistry, as described. The sample was loaded onto a single Sequel II SMRT cell (v3.2 chemistry) and sequenced by the University of Washington PacBio Sequencing Services. The full composition and sample barcode IDs of the multiplexed library are listed in **Table S5**.

Plasmid DNA was prepared in a separate multiplexed library. Plasmids were linearized with KpnI prior to multiplexed library preparation.

### Quantification of m6A/A by UHPLC-MS/MS

Samples for quantification were treated as previously described with minor modifications (Kong *et al*., 2022). In brief, 30-50 ng of DNA from each sample was mixed with 0.02 U phosphodiesterase I (Worthington, cat# LS003926), 1 U Benzonase (Millipore Sigma, cat# E1014), and 2 U Quick CIP (NEB, cat#M0525S) in digestion buffer (10 mM Tris, 1 mM MgCl, pH 8 at RT) for 3 hours at 37°C. Single nucleotides were separated from the enzymes by collecting the flow-through of a Nanosep centrifugal filter (MWCO 3 kDa, Pall, cat#OD003C33). The UHPLC-MS/MS analysis of adenosine and m6A was performed on an ACQUITY Premier UPLC System coupled with XEVO-TQ-XS triple quadrupole mass spectrometer. UPLC was performed on a ZORBAX Eclipse Plus C18 column (2.1 × 50 mm I.D., 1.8 μm particle size) (Agilent, cat# 959757-902) using 10-90% linear gradient of solvent B (0.1% acetic acid in 100% methanol) in solvent A (0.1% acetic acid in water) within 4 minutes and a flow rate of 0.3 ml/min. MS/MS analysis was operated in positive ionization mode with 3000 V capillary voltage as well as 150°C and 1000 L/Hour nitrogen drying gas. A multiple reaction monitoring (MRM) mode was adopted with the following m/z transition: 252.10 −>136.09 for dA (collision energy, 14 eV), and 266.2->150.2 for m6A (collision energy, 15 eV). MassLynX was used to quantify the data.

A calibration curve was generated with 11 mixtures containing different ratios of 2-deoxyadenosine (>99%, FisherScientific, cat#AAJ6388606)(A) to N6-methyl-2-deoxyadenine (>99%, FisherScientific, cat# AAJ64961MD)(m6A). A new standard was measured and used for each run. The standard was fit to a third-degree polynomial (equation 1) with y as m6A percentage (%) and x as the quantified MS peak area of m6A over the sum of adenosine and m6A peak area.

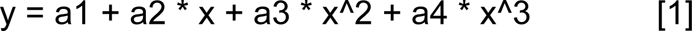

## Acknowledgments

The authors thank Sayeh Gorjifard for organizing the University of Washington Genome Sciences Hackathon, which initiated this work, Michelle Noyes for designing the fibertools logo art, and Tonia Brown for assistance in editing this manuscript.

## Author contributions

Conceptualization and design: M.R.V., A.J., S.C.B., and A.B.S. Experimental design and execution: S.C.B., Y.M., J.R., B.J.M., and A.B.S. Preliminary implementation: M.R.V., A.J., S.C.B., A.T.M., M.O.H., E.S., C.F., and T.L. Final implementation: M.R.V. and A.J. Supplemental material organization: M.R.V., S.C.B., and A.J. Display items: M.R.V., A.J., S.C.B., and A.B.S. Manuscript writing: M.R.V., S.C.B., and A.B.S. with input from all authors.

## Data and materials availability

All K562 and CHM1 PacBio data was generated as part of a previous study and is available at the following GEO accession: GSE226394. All additional PacBio generated as part of this study is available through the SRA under Bioproject ID: PRJNA956114. Cell lines obtained from the NIGMS Human Genetic Cell Repository at the Coriell Institute for Medical Research include GM12878 and K562. Additional data used in analysis is made available on Zenodo (https://zenodo.org/record/7809229).

## Code availability

Code for training (https://github.com/fiberseq/train-m6A-calling), evaluation (https://github.com/mrvollger/Fiber-m6A-figures-and-tables), and execution (https://github.com/fiberseq/fibertools-rs) of fibertools is publicly available on GitHub.

