## supplemental information for "DNA-m6A calling and integrated long-read epigenetic and genetic analysis with fibertools"

#### Supplemental Methods

##### Fibertools algorithm

We derive m6A labels from GMM-filtered ipdRatios from ipdSummary. These labels can contain false positives. The lack of clean m6A labels makes the supervised training approach less suitable since it assumes that accurate labels are available. Therefore, we developed a semi-supervised approach, which assumes that our m6A class is a mixed population of true and false positives and that our non-m6A class is a clean set. Our training approach derives from Percolator and mokapot, proteomics tools for identifying peptides from mass spectrometry data (Käll *et al.*, 2007; Fondrie and Noble, 2021).

Given a set of candidate m6A calls, Fibertools aims to maximize the number of m6A calls at a target estimated precision. First, we split our dataset into training and validation sets stratified by class labels (see **Table S3** for the training and validation datasets sizes for different Fiber-seq chemistry experiments). Then our method proceeds in two phases. In the first phase, we use the central base's interpulse duration (IPD) score as a classifier and generate an m6A classification score for all examples in the validation set. The validation examples are then ranked by the classification score, and estimated precision is computed at every score threshold. In general, precision is defined as the ratio of true positives over the total number of positive calls (i.e., true positives plus false positives). In our case, for a set of m6A ($P$) and non-m6A calls (*N*) meeting a score threshold, the denominator of the precision calculation is simply *P* + *N*. For the numerator, i.e., the number of true positives, we assume that some of the *P* m6A calls are false positives. In particular, we assume that each non-m6A call above the threshold corresponds to *c* false positive m6A calls, where *c* is the ratio of positive to negative labels in our validation dataset. Thus, we define estimated precision, *EP*, as

$$\frac{P - cN}{P + N}$$

Using a ranked list of *EP* at different score thresholds, we select the score threshold with the target *EP* in the validation set. At the end of this phase, we have a score threshold for identifying m6A calls at the target estimated precision. In practice, 12-14% of our data had m6A labels, and we used the target estimated precision of 95%.

The second phase is iterative, and each iteration consists of three steps. The first step is selecting a high-confidence m6A training set using the score threshold from the first phase for the first iteration and step 3 of the second phase for subsequent iterations. The second step consists of training a CNN model on this training data. In the final step, the validation data is rescored using the trained CNN model from the second step, and a new score threshold is generated with the rescored validation data. We will now describe each step in detail.

In the first step, a score threshold is used to select a putative m6A set. This score threshold is derived from the IPD score classifier in the first iteration and a trained CNN in subsequent iterations. At the end of this step, we have a training set selected using a given score threshold and classifier.

The second step is training a CNN model to discriminate between m6A and non-m6A examples. We initialize our model using the CNN model from the supervised approach. This transfer learning approach allows fast convergence of our training procedure. We retain the same training hyper-parameters as the supervised approach. At the end of this step, we produce a CNN classifier trained on training data from the previous step. We train the 2.2, 3.2, and Revio chemistry models for two epochs. During an epoch, we compute average precision on the validation data every x iteration, where x refers to the number of batches corresponding to 10% of training data. We save the model with the best average precision on validation data. Training for two epochs was sufficient to learn a model with better average precision than the previous round for all three Fiber-seq chemistries.

The final step is finding a new score threshold by rescoring the validation examples using the trained CNN from step two. The initial process of estimating *EP* at different score thresholds is repeated, and a new score threshold with a target *EP* (95%) is selected.

In the case of a successful second phase training, the number of positives in the validation data identified at target precision increases with every iteration and plateaus when most m6A examples in the validation data have been identified. We define two conditions for convergence, both of which must be satisfied. First, more than 70% of putative m6A calls from the validation set have been identified. Second, the number of additional m6A calls in a new iteration is less than 1% of the total putative m6A calls. In practice, it took 12, 11, and 3 repetitions of phase two training to converge 2.2, 3.2, and Revio chemistry Fiber-seq experiments, respectively (**Fig. S9**).

**Algorithm 1 The Fibertools Algorithm** The input variables are: $\mathcal{TD}$ = training data, $\mathcal{VD}$ = validation data, $c$ = ratio of positive and negative m6A calls in $\mathcal{TD}$ and $\mathcal{VD}$, $t$ = target estimated precision threshold, *IPDClassifier* = m6A classifier using the IPD score of the central base, $\mathcal{I}$ = the number of iterations.

```
procedure Fibertools(TD, VD, c, t, IPDClassifier, I)
    s_t ← scoreAtTargetPrecision(IPDClassifier, VD, t, c)      ▷ Compute score threshold s_t at t
    for i ← 1...I do
        TD_t ← selectTrainingSet(TD, s_t)                      ▷ Compute new training set TD_t
        CNN_w ← trainCNN(TD_t)                                 ▷ Train the CNN classifier CNN_w
        s_t ← scoreAtTargetPrecision(CNN_w, VD, t, c)          ▷ Recompute s_t
    end for
    return (s_t, CNN_w)
end procedure
```

### Supplemental Figures

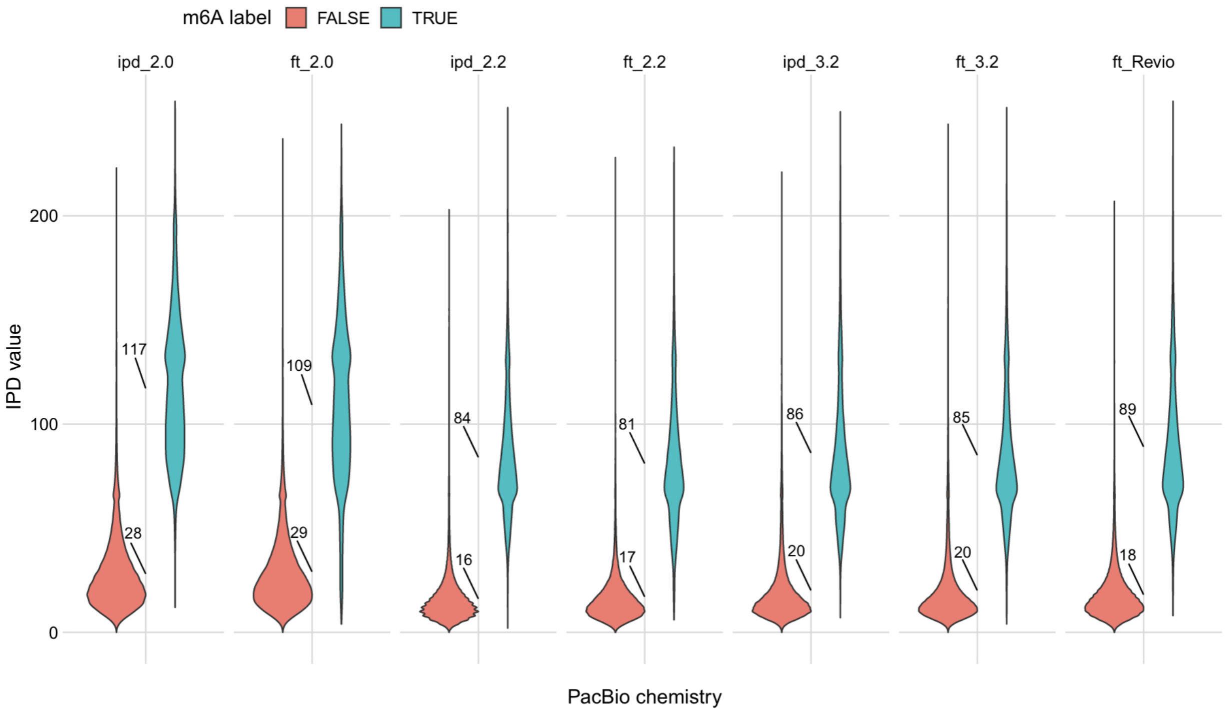


**Figure S1. Interpulse duration (IPD) at m6A and adenine base pairs.** Violin plots of the IPD value following unmethylated adenines (FALSE, red) and m6As (TRUE, cyan). Different columns show values for ipdSummary (ipd) and fibertools (ft) calls and different chemistries (2.0, 2.2, 3.2, and Revio).

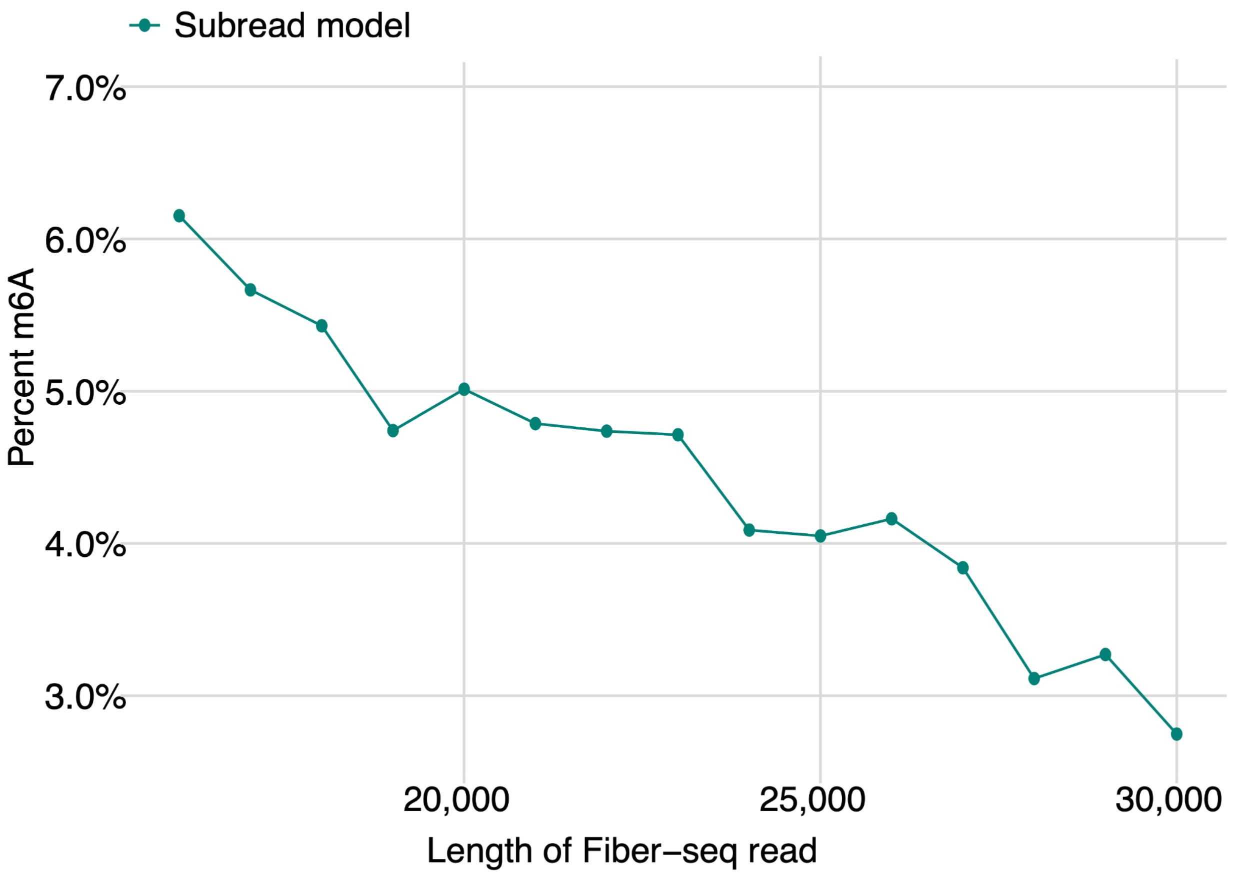


**Figure S2. Percent m6A as a function of read length.** Percent of methylated adenines relative to all adenines called by the subread model as a function of read length (2.2 chemistry model).

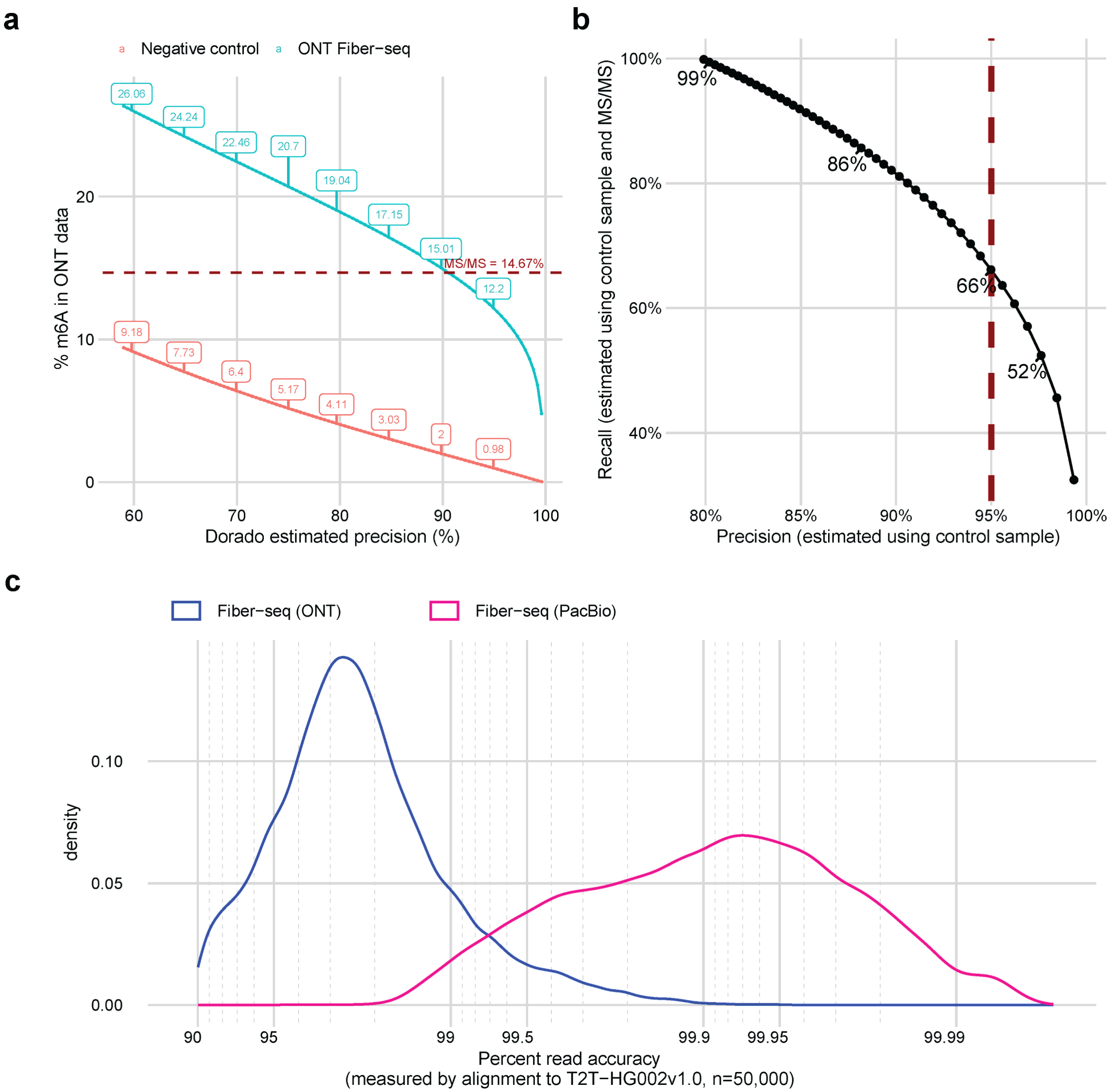


**Figure S3. Dorado m6A calling on ONT Fiber-seq. a)** Comparison of the percent m6A observed in ONT Fiber-seq (blue) and a gDNA negative control sample (red) as a function of the precision reported by Dorado. The horizontal line indicates the percent m6A in the ONT Fiber-seq sample as measured by MS/MS. **b)** Estimated precision-recall curve for Dorado m6A calling. The text indicates the estimated recall at 10% increments of precision, and the red line marks a 95% precision level. We estimated the fraction of true positive calls by subtracting the fraction of m6A calls in the gDNA negative control from the fraction of m6A calls in ONT Fiber-seq. We calculated precision by dividing the estimated fraction of true positive calls over the total fraction of m6A calls, and we calculated recall by dividing the estimated fraction of true positive calls by the total percent m6A in the ONT Fiber-seq sample (14.67%) which we identified using MS/MS. **c)** Distribution of read accuracy for PacBio (pink) and ONT (blue) Fiber-seq HG002 samples, as determined by comparison to the reference genome.

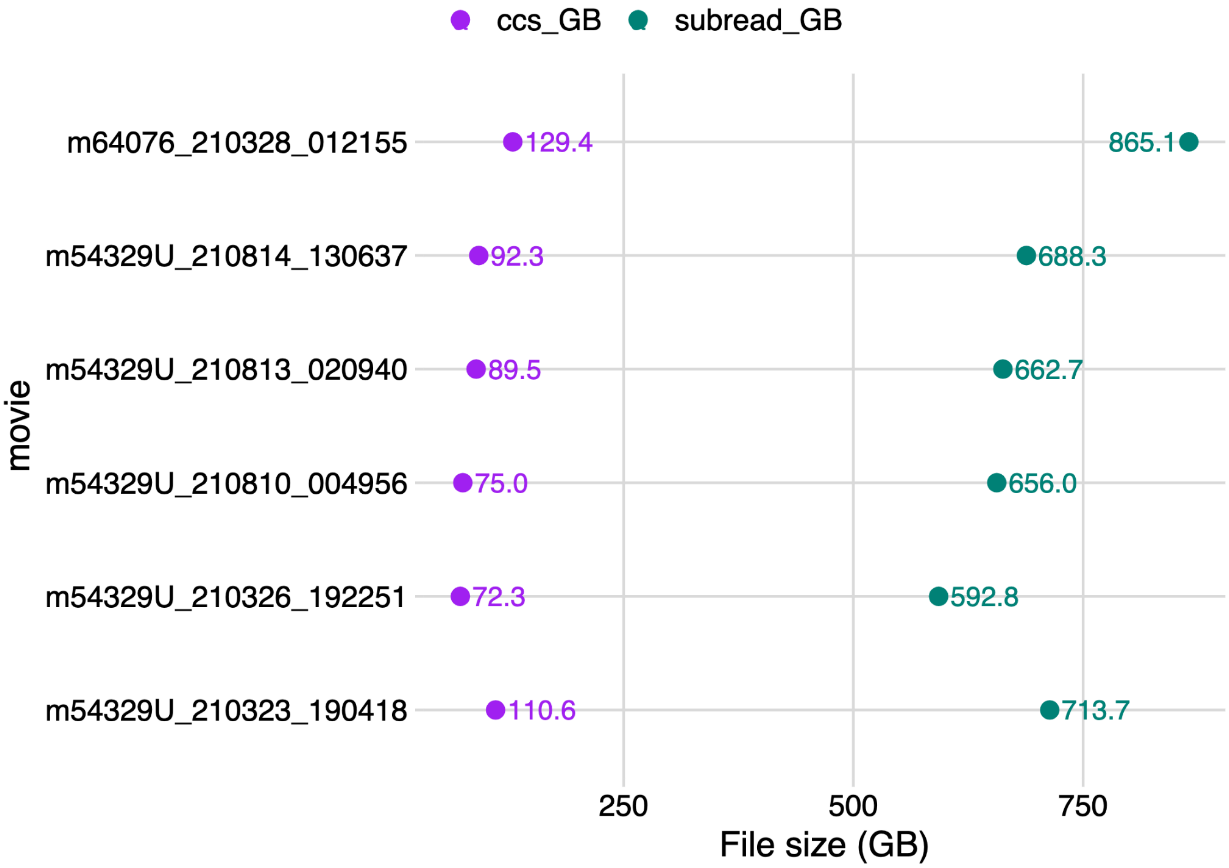


**Figure S4. Size comparison of CCS and subread files.** File size in GB for CCS files with kinetics (purple) compared to the equivalent subread file (green).

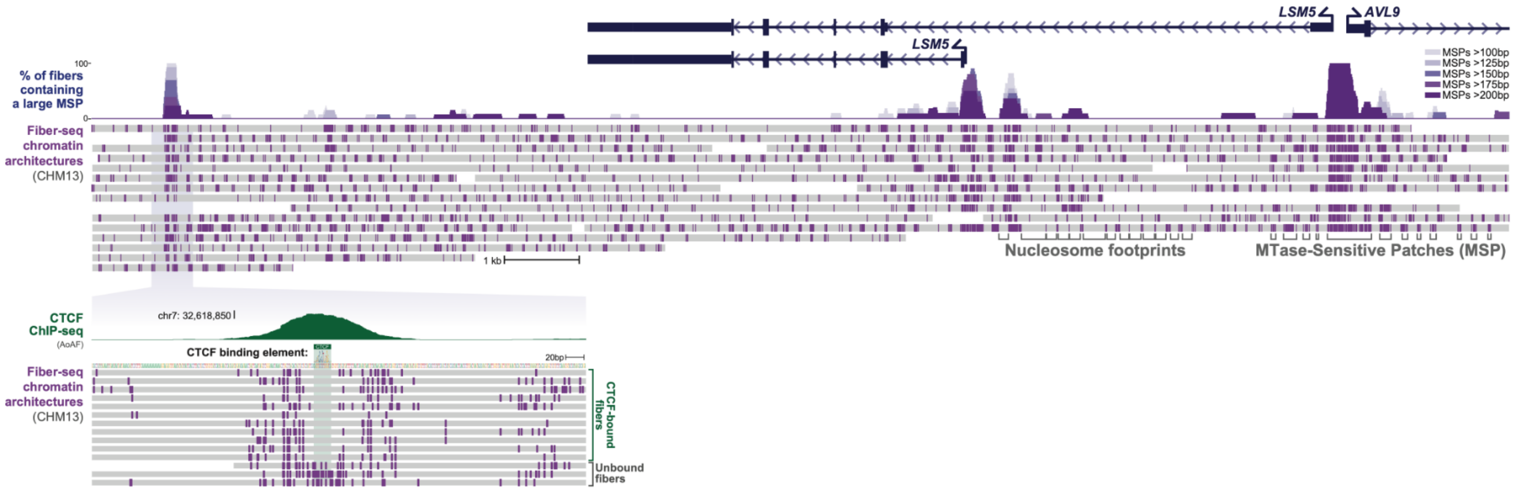


**Figure S5. Example locus showing Fiber-seq data in CHM13.** Each line represents an individual sequencing read. Purple ticks represent m6A events called by the subread model. Nucleosome footprints (~150 bp patches devoid of m6A) and MTase-sensitive patches (internucleosomal region at least 65 bp) are indicated. The inset is a zoomed-in view of a CTCF binding element depicting fibers that are bound and unbound (bottom three reads) by CTCF.

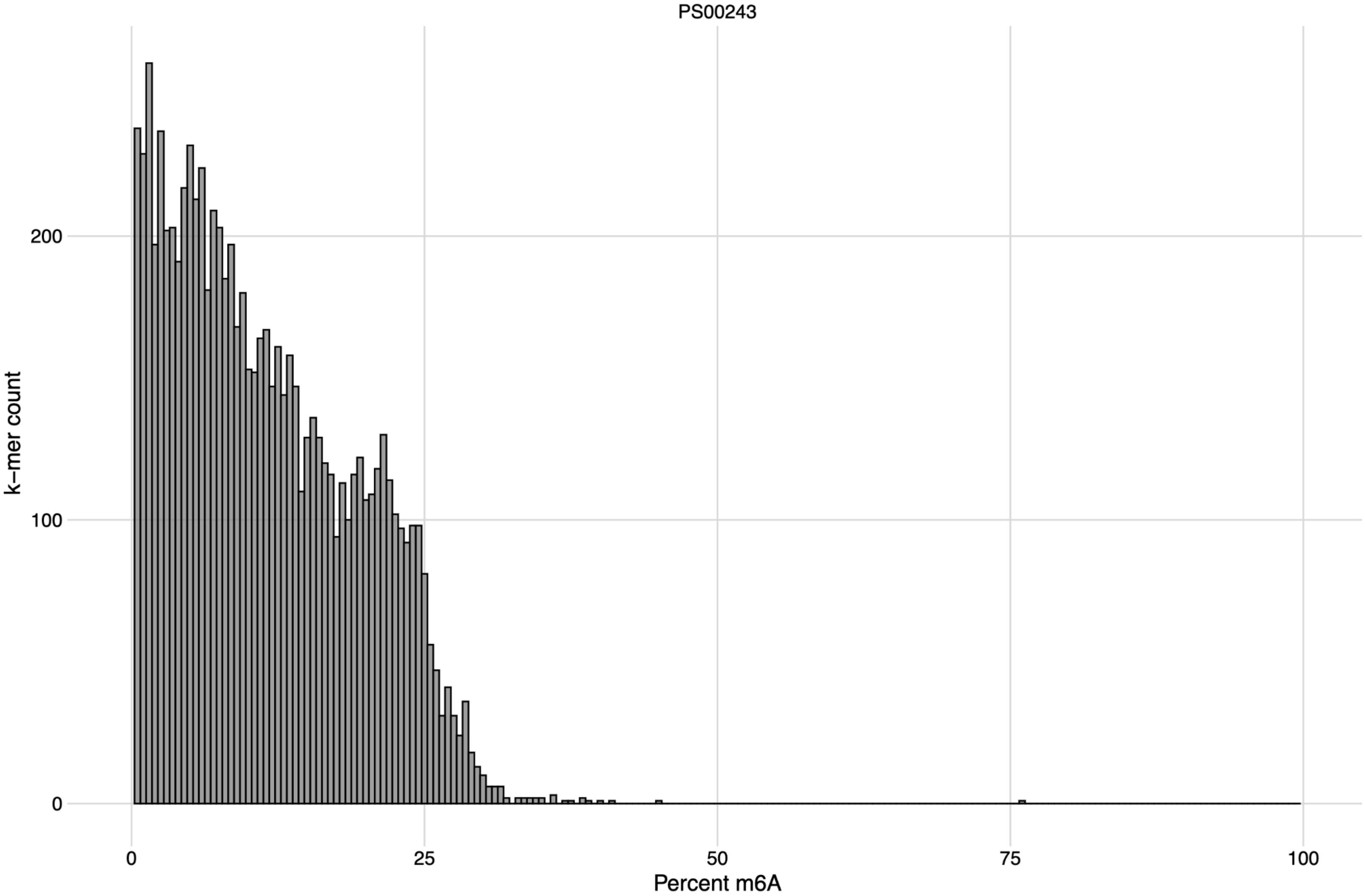


**Figure S6. Percent m6A of all different heptamers.** Count of 7-mers (y-axis) binned by % of that 7-mer containing a central m6A with respect to all instances of the same 7-mer. m6A calls were made by the fibertools 3.2 model on Fiber-seq K562 data.

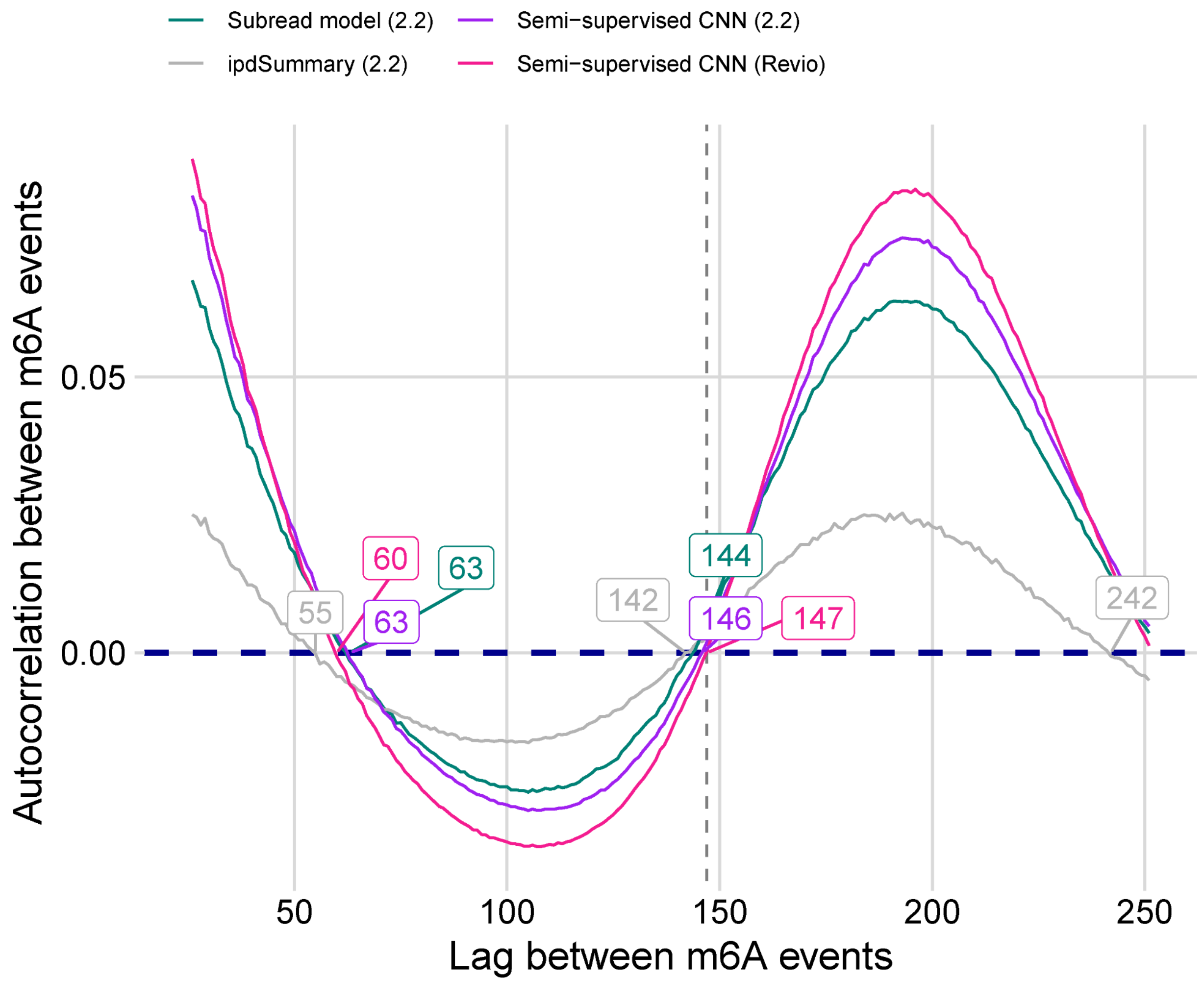


**Figure S7. m6A autocorrelation from different callers.** Autocorrelation is shown for calls made by the subread model (teal), ipdSummary model (gray), semi-supervised CNN for PacBio 2.2 chemistry (purple), and semi-supervised CNN for PacBio Revio chemistry (pink).

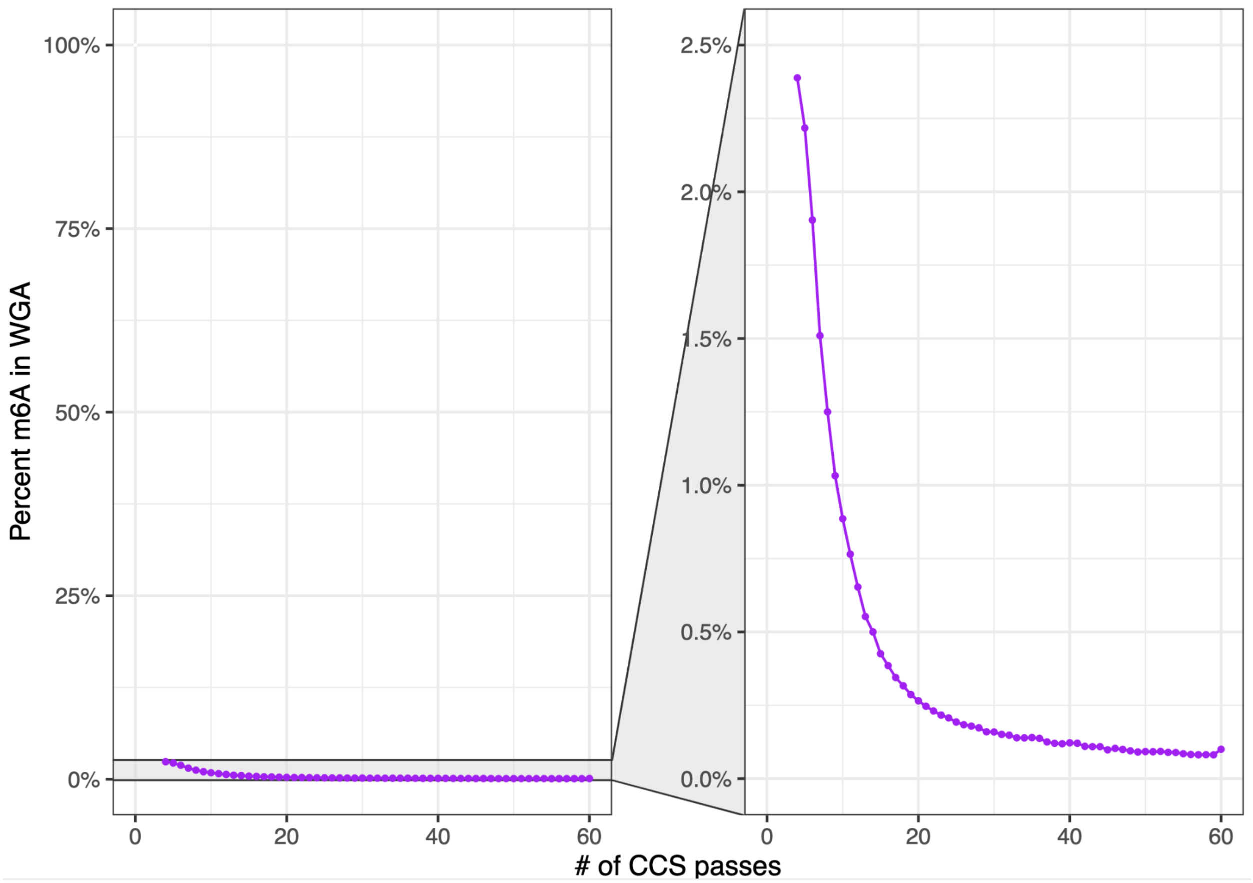


**Figure S8. False positive rate in WGA data as a function of CCS passes.** Percentage of m6A calls with respect to all adenines as a function of CCS pass number.

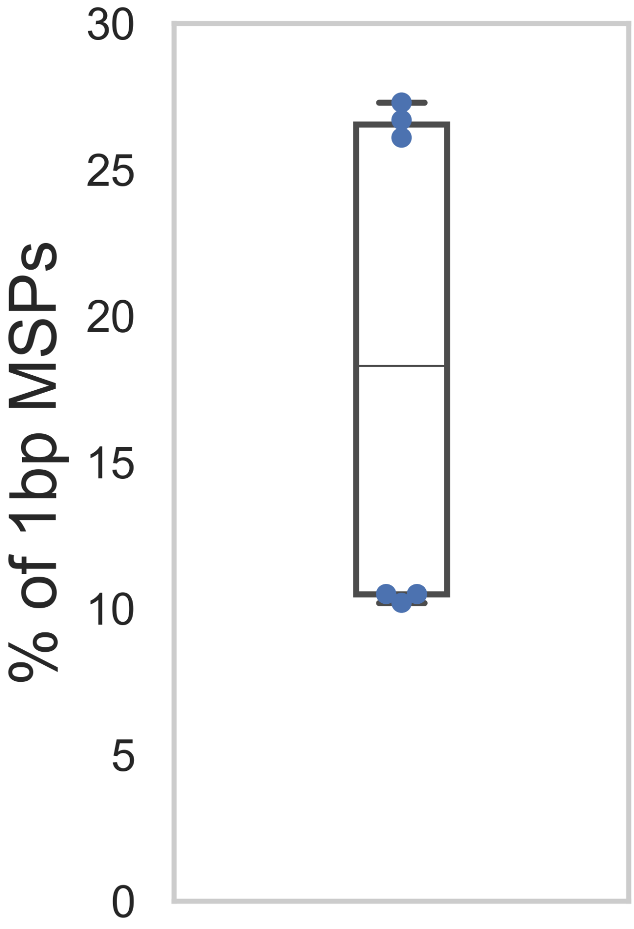


**Figure S9: Single m6A internucleosomal linker regions.** Percent of all internucleosomal linker regions defined by only a single m6A separating two nucleosomes. Data is shown for three SMRT cells PS00075 and three SMRT cells of PS00109 (n=6).

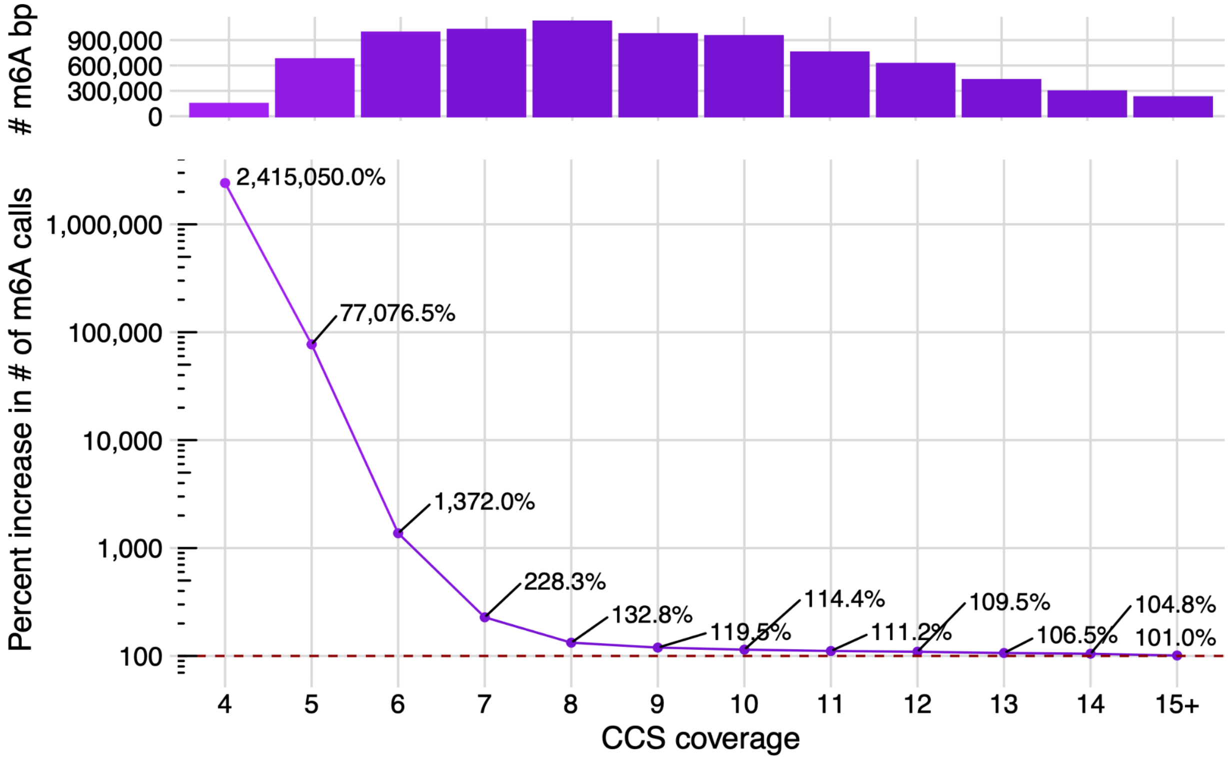


**Figure S10. Increase in m6A calls for fibertools.** The top plot shows the number of m6A bases in reads with increasing CCS coverage. The bottom plot shows the percent increase of m6A calls for fibertools over the subread model as a function of CCS coverage. The x-axis is shared between the top and bottom plots.

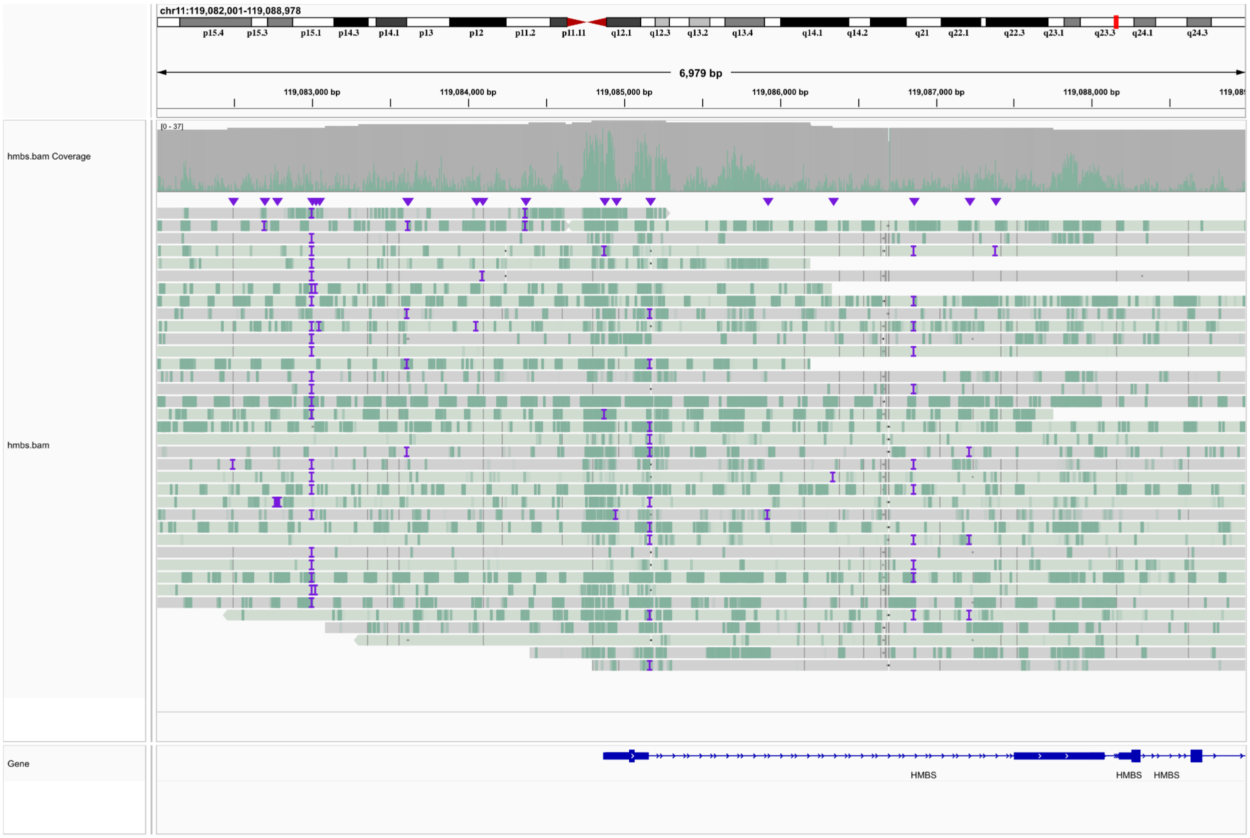


**Figure S11. IGV screenshot of the *HMBS* promoter.** Screenshot shows the encoding of m6A methylation (green) in the BAM format using the MM and ML tags for the *HMBS* promoter with Fiber-seq data from K562.

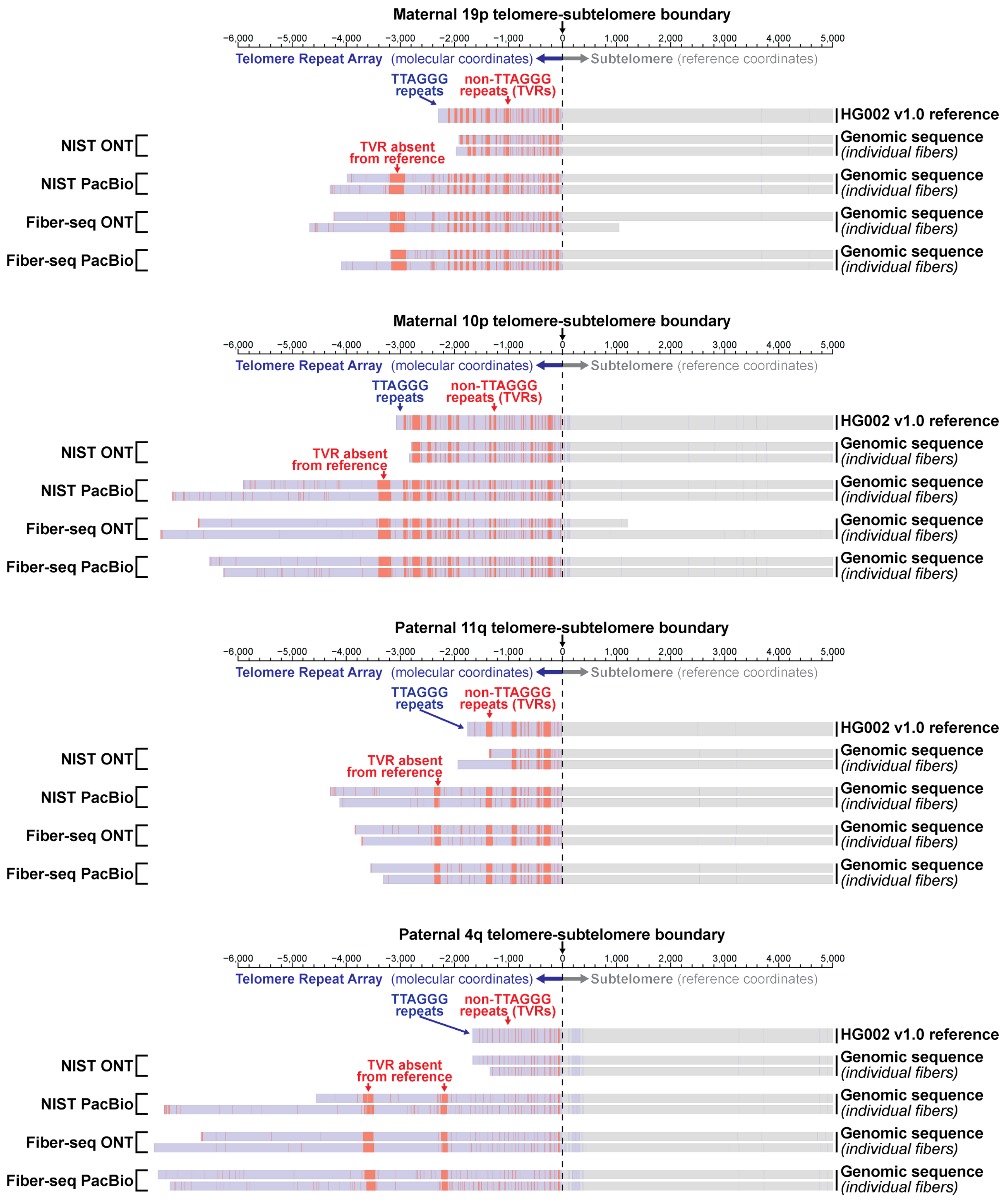


**Fig. S12. Multiple non-TTAGGG telomere variant repeats (TVRs) (Baird *et al.*, 1995; Allshire *et al.*, 1989) are absent from the HG002 reference sequence.**

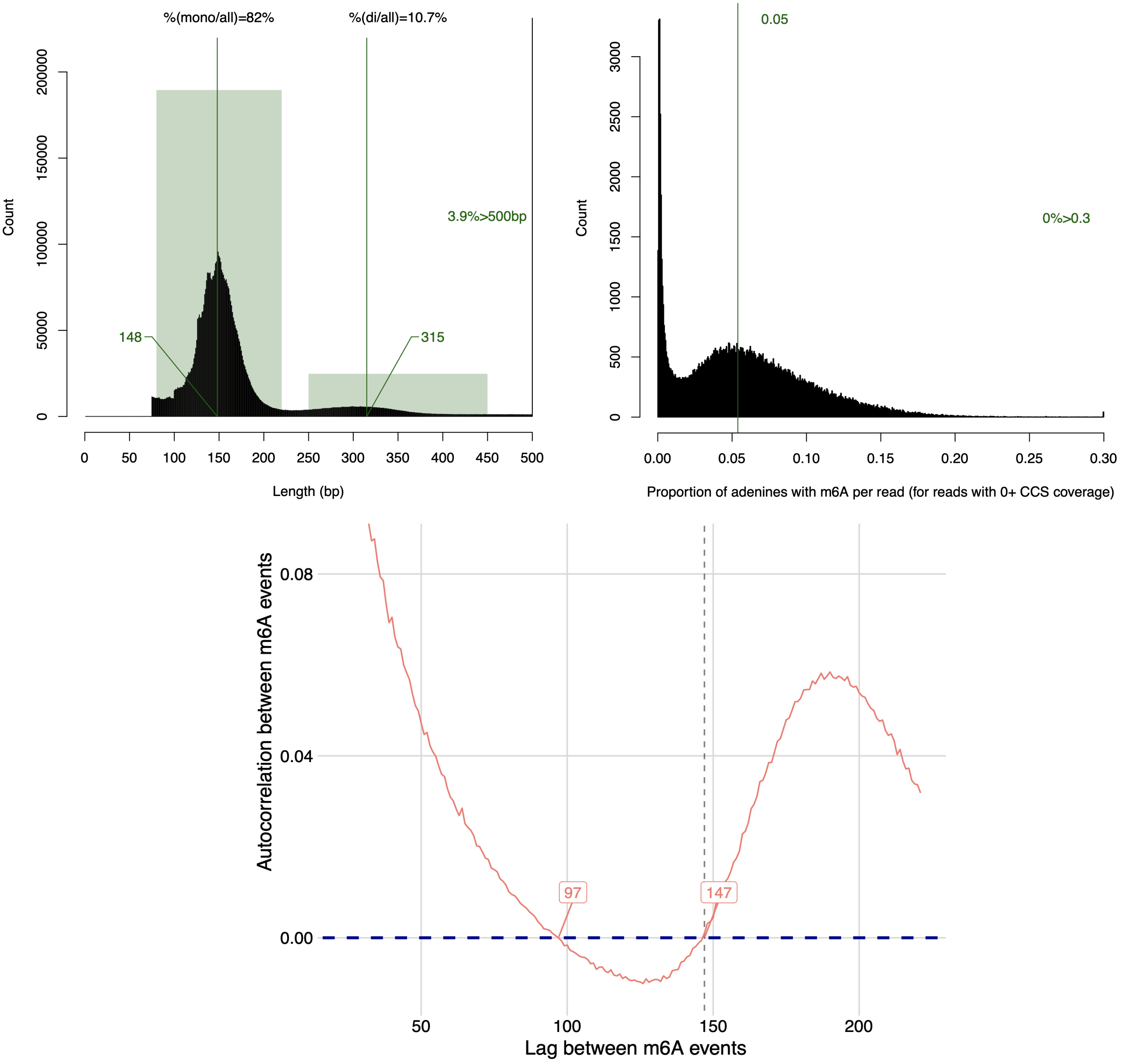


**Figure S13: Fibertools applied to a sample prepared with EcoGII.** *top left)* Histogram of nucleosome lengths within the EcoGII sample. *top right)* Histogram of proportion of adanines with m6A in the sequencing reads. *bottom)* Autocorrelation between adjacent m6A calls, identifying the exact length of the nucelosomes within the data (147 bp).

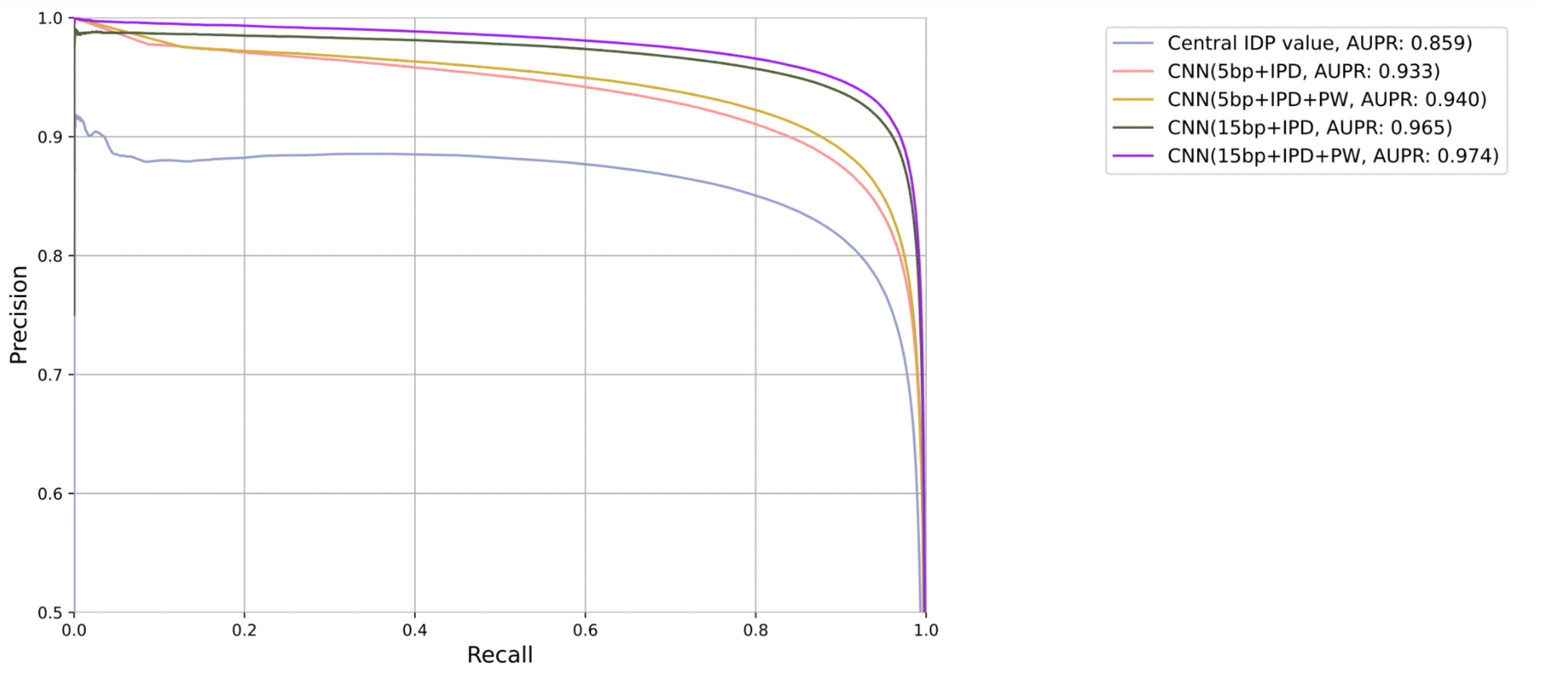


**Figure S14. Ablation study of the full-supervised CNNFibertools.** Precision-Recall curve for five versions of the fibertools fully-supervised CNN model. The Area Under the Precision-Recall (AUPR) curve values of each variant are reported in the legend. A description of each variant follows. *Central IPD value*: A method that uses only the IPD value of the central adenine base without any surrounding sequence information, IPD values, or Pulse Width values. *CNN 5bp+IPD*: A CNN model using IPD values and sequence information from a 5 bp window surrounding the central adenine base. *CNN 5bp+IPD+PW*: A CNN model using IPD values, Pulse Width values, and sequence information from a 5 bp window surrounding the central adenine base. *CNN 15bp+IPD*: A CNN model using IPD values and sequence information from a 15 bp window surrounding the central adenine base. *CNN 15bp+IPD+PW*: A CNN model using IPD values, Pulse Width values, and sequence information from a 15 bp window surrounding the central adenine base.

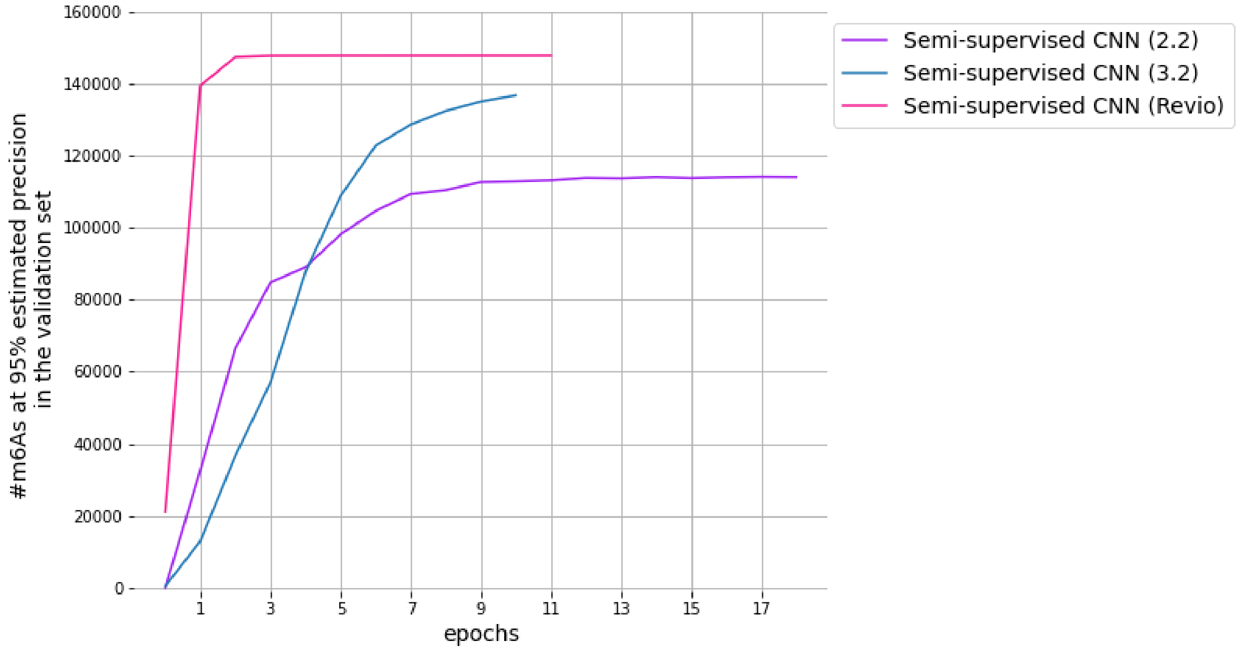


**Figure S15: Number of m6A calls identified at 95% precision increases with semi-supervised training epoch.** The number of m6A calls identified at 95% estimated precision in the validation data from three different chemistries (2.2, 3.2, and Revio) as the training epochs progress. The x-axis shows the number of training epochs, and the y-axis shows the number of m6A calls in the validation set at 95% estimated precision (see Supplementary Methods for the definition of estimated precision).
